# Fast and Powerful Genome Wide Association Analysis of Dense Genetic Data with High Dimensional Imaging Phenotypes

**DOI:** 10.1101/179150

**Authors:** Habib Ganjgahi, Anderson M. Winkler, David C. Glahn, John Blangero, Brian Donohue, Peter Kochunov, Thomas E. Nichols

## Abstract

Genome wide association (GWA) analysis of brain imaging phenotypes can advance our understanding of the genetic basis of normal and disorder-related variation in the brain. GWA approaches typically use linear mixed effect models to account for non-independence amongst subjects due to factors such as family relatedness and population structure. The use of these models with high-dimensional imaging phenotypes presents enormous challenges in terms of computational intensity and the need to account multiple testing in both the imaging and genetic domain. Here we present method that makes mixed models practical with high-dimensional traits by a combination of a transformation applied to the data and model, and the use of a non-iterative variance component estimator. With such speed enhancements permutation tests are feasible, which allows inference on powerful spatial tests like the cluster size statistic.

## Introduction

Genome-wide association studies (GWAS) of neuroimaging data can advance our understanding of human brain by discovering genetic variants associated with normal and disorder-related phenotypic variance in brain structure and function^1–6^. The genetic association analysis with the quantitative phenotypes from structural (i.e. brain volume, cortical thickness, white matter integrity) or functional imaging modalities (brain response to particular cognitive task or resting state) at hundred thousand locations in the human brain present statistical challenges including statistical power, multiple comparisons correction and like other association studies correction for population structure, a term that encompasses cryptic/family relatedness and population stratification.

In the GWA studies of unrelated individuals, non-independence due to latent population stratification or due to unknown (often termed cryptic) relatedness^7, 8^ is generally thought to be a confounding factor that can lead to excessive false positives when ignored. This type non-independence has been studied throughly in the recent GWA era^9–13^. While genomic data can be used to control for population stratification by including the top principal components as a fixed effect covariates in a linear regression model^14^, usually individuals with close estimated relatedness from identity-by-state (IBS) matrix or different ethnicities are excluded from the study sample. This might not be a problem in genetic studies with 4 digits sample sizes, but may make substantial differences in GWA studies with neuroimaging phenotypes where sample size is much smaller. Also, even in a carefully design GWA study, it is hard to avoid spurious associations because of population structure; in particular it is likely that in studies with large sample sizes i.e the UK biobank some level of population structure are induced within a same population. Moreover, GWA studies with neuroimaging phenotypes require fitting a marginal model at each point (voxel/element) in the brain, the large number of measurements presents a challenge both in terms of computational intensity and the need to account for elevated false positive risk because of the multiple testing problems both in terms of number of elements in image and number of markers being tested. Although the emergence of large scale neuroimaging consortia like ENIGMA or CHARGE can help to conduct well-powered genetic association studies through meta analysis framework, still it is crucial to use a powerful statistical method at the site level. Hence, there is a compelling need for a analytical technique that addresses these challenges.

There has been great interest in the field of quantitative genetic to develop sophisticated statistical methods to control population structure in GWA studies of unrelated individuals. Linear mixed models (LMM) allowing for the rigorous testing of genetic associations (and, more generally, fixed effects) have long been employed in human genetics as the standard to exploit and/or correct for the non-independence among subjects due to known familial relatedness in pedigree-based studies^15–19^. Linear mixed effect models using molecularly-derived empirical relatedness measures have gained popularity recently for both studies of related and unrelated individuals since they do not require prior information on biological relatedness and/or represent a framework where such complexities are automatically accounted for. One popular utilization of the LMM is as an alternative method to linear regression models where the association statistic incorporates a component of trait variance that is explained by a genetic relationship matrix (GRM) that captures the genome-wide similarity between “unrelated” individuals by modeling it as a random effect^20–31^. Additionally, it has been shown that the correction for the problem of latent population structure in GWA with an LMM is both effective and power preserving^30–32^.

Due to the required inversion of potentially large matrices, the general LMM is computationally intensive where the complexity includes the deriving of the GRM, variance component parameter estimation, fixed effect estimation, and the calculation of the required association statistic for each marker grows with sample size and number of candidate markers for association testing. Several approximate or exact methods have been proposed to speed up LMM-based testing. Approximate methods assume the total polygenic random effect is same for all markers under the null hypothesis of no marker effect, hence the relevant residual genetic variance component is estimated only once using all markers. In contrast, exact methods, which is the correct LMM practice^24, 25, 30, 30^, estimate a residual variance component conditional on each marker’s effect. In studies of “unrelated” subjects, this residual variance component often involves re-estimation of the GRM which is constructed excluding the candidate marker and surrounding markers in linkage disequilibrium.

The LMM efficiency in controlling confounding factors in the genetic association analysis and possible boost in power inspires using it with high-dimensional imaging phenotypes. However fitting LMM at each voxel/ROI in the brain is computationally intensive or even intractable at the voxel level while variance component estimation relies on likelihood function optimisation using numerical methods. Moreover, search for genetic association across the genome at different locations with imaging phenotypes requires intense multiple testing corrections both for number of elements in an image and number of markers. Whether the association analysis is conducted at the reduced search space in the brain i.e., summary measure from a region of interest or voxel level, naive application of bonferroni correction for number of hypothesis testing in the image with usual GWA P-value leads to invalid statistical inference procedure while it ignores complex spatial dependence between elements in the imaging phenotypes. The parametric null distribution of cluster size^34^ or threshold free cluster enhancement (TFCE) statistics^35^ that are the most common and sensitive inference tools in imaging, could be invalid due to untenable stationary assumption^36, 37^ or in the later case be unknown. Familywise error rate (FWE) correction, controlling the chance of one or more false positives across the whole set (family) of tests^38^ requires the distribution the maximum statistic, can be computed for either voxels/ROI or cluster size with permutation test^39^ which is the standard tool to conduct inference in neuroimaging.

Despite many analytical techniques have been developed to accelerate the GWA with LMM, these advances do not eliminate problems related to numerical optimisation nor multiple testing problem. This paper makes two major contributions to reduce the complexity of LMM in the genetic association specifically with the imaging phenotypes. First, variance component estimation step computational cost is reduced using non-iterative one-step random effect estimator^40^. Second, complexity of association testing is dramatically decreased with projecting the model and phenotype to a lower dimension space.

To our knowledge the fastest implementation of exact LMM is Fast-LMM^24^, which transforms the phenotype and LMM model with the genetic similarity matrix (GRM) eigenvectors and uses a profile likelihood approach to simplify variance component estimation. The eigenvector matrix diagonalisation along with the profile likelihood with only one variance parameter reduces optimisation time substantially. In Fast-LMM the covariance matrix is estimated only under the null hypothesis of no marker effect, and then a generalized least squares (GLS) is applied to estimate the marker effect and the likelihood ratio test is used for hypothesis testing. Note that small sample size behavior of this approach has not been validated, using it for association analysis of imaging phenotypes with only, say 300, subjects might not be valid. In addition to concerns about the finite sample validity, Fast-LMM requires numerical optimisation for each element (voxel/ROI) of image that makes it computationally intensive or essentially impractical for large-scale imaging phenotypes.

The key to our method is the projection of the phenotype and the model to a lower dimension space, and a score statistic for association testing. This projection is based on the eigenvectors of the adjusted GRM for the fixed effect nuisance terms. In this setting, the projected phenotype likelihood function is equivalent to that used with restricted maximum likelihood (REML) of the LMM (**??**), going forward we call this approach *simplified REML*. While both models have the same statistical properties, our particular projection provides several computational benefits that reduces LMM complexity dramatically as follows: (I) As we described in previous work^40^, the diagonalized covariance allows a non-iterative one-step variance component estimator, taking the form of a weighted regression of squared projected data on eigenvalues of the GRM adjusted for nuisance fixed effect terms, an approach that we henceforth call *WLS-REML* (weighted least squares REML); (II) The regression form of our estimator is easily vectorized, meaning that many image elements and SNPs can be tested in a single and fast computational test in several high-level programming languages (Online Methods); (III) Finally, the simplicity and fast computation of the score test statistic makes permutation testing feasible, allowing exact, non-parametric control over the FWE, accounting for the number of tests conducted over all image elements and markers. Two permutation schemes can be defined, free and constrained, where in the latter case the permutation is confined to exchangeability blocks defined based on the eigenvalues distribution.

The reduced computational complexity of our method represents a significant advance over existing methods. The complexity of LMM association has two components, one for the variance component estimation, the other is for fixed effect parameter estimation and test statistic computation. For a GWA over *S* markers and *V* imaging phenotype elements on *N* individuals, the variance component likelihood optimization complexity of FaST-LMM is *O*(*N*^3^ + *INV*), where *I* is the average number of iterations, while for WLS-REML the random effect estimator (Online Methods) it is *O*(*N*^3^ + *NV*) (the common *O*(*N*^3^) term is the time complexity of the GRM eigendecomposition). More critically, the estimation and test statistic computation complexity of FaST-LMM is *O*(*SPN*^2^ *V*), where *P* is the number of nuisance fixed effects, while for WLS-REML (Eq. (15)) this is *O*(*SNV*), a substantial reduction for imaging phenotypes when number of image elements *V* is much bigger than the sample size *N*. Even for a single trait GWA (*V* = 1), our proposed projection reduces the association (Eq. (12)) complexity to *O*(*SN*) which is significantly less than FaST-LMM for large sample GWA.

In our previous work^40^ we introduced WLS-ML random effect estimator that exploits this one-step optimization approach combined with eigen-rotation of phenotype and model (see Online Methods for more details). The non-iterative estimator has a simple form, with variance components estimated as a weighted regression of squared ordinary least square residuals on eigenvalues of the GRM, and fixed parameters estimated with a weighted regression of eigen-transformed phenotype on eigen-transformed model. In this paper we evaluate our non-iterative ML and REML estimators (WLS-ML and WLS-REML) with their fully converged counterparts (Full ML or Full REML), comparing score, likelihood ratio (LRT) and Wald tests on intensive simulation studies. The score test based on the simplified REML function is compared with FaST-LMM using the simulation study and real data analysis.

## Results

### Simulation Results

Simulation results on the accuracy of genetic random effect 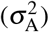 estimation shows that the non-iterative one-step approaches are similar to their fully converged counterparts (Supplementary Figure 1), using either likelihood or restricted likelihood functions. When the data are independent 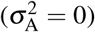, the methods are indistinguishable in terms of bias and mean squared error (MSE). When 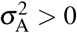, the fully converged methods have less bias, but the difference is modest in absolute value; in terms of MSE, the non-iterative one-step methods have just slightly worse performance. The first simulation also shows good performance of fixed effect (*β*_1_) estimation (Supplementary Figure 2). Both the non-iterative one-step and fully converged have similar bias and MSE, with WLS-REML again closely following fully converged REML.

Simulations show that the false positive rates for the fixed effect score test for *H*_0_: *β*_1_ = 0 (Supplementary Figure 3a) are nominal; for both simplified ML or REML functions, for all simulation settings considered, test statistic type and type of random effect estimator, the false positive rates lay within the Monte Carlo confidence interval (MCCI) (See also Supplementary Figures 4a & 4b).

The simulation results on the power of score test reveal negligible differences between the random effect estimation methods (Supplementary Figure 3b). Similar findings are obtained for the power of LRT and Wald tests (Supplementary Figures 4c & 4d). Like the parametric approach, we found that both permutation schemes, free or permutation within exchangeability blocks, control the false positives at the nominal level 5% (Figure 1a and Supplementary Figures 5a & 5b), and could provide nearly equivalent power (Figure 1b, Supplementary Figures 5c & 5d) for all statistics either based on the simplified ML or REML functions. However, for all test statistics and 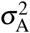, the free permutation scheme is slightly more powerful than the constrained permutation test when a kinship matrix is used.

**Figure 1.**
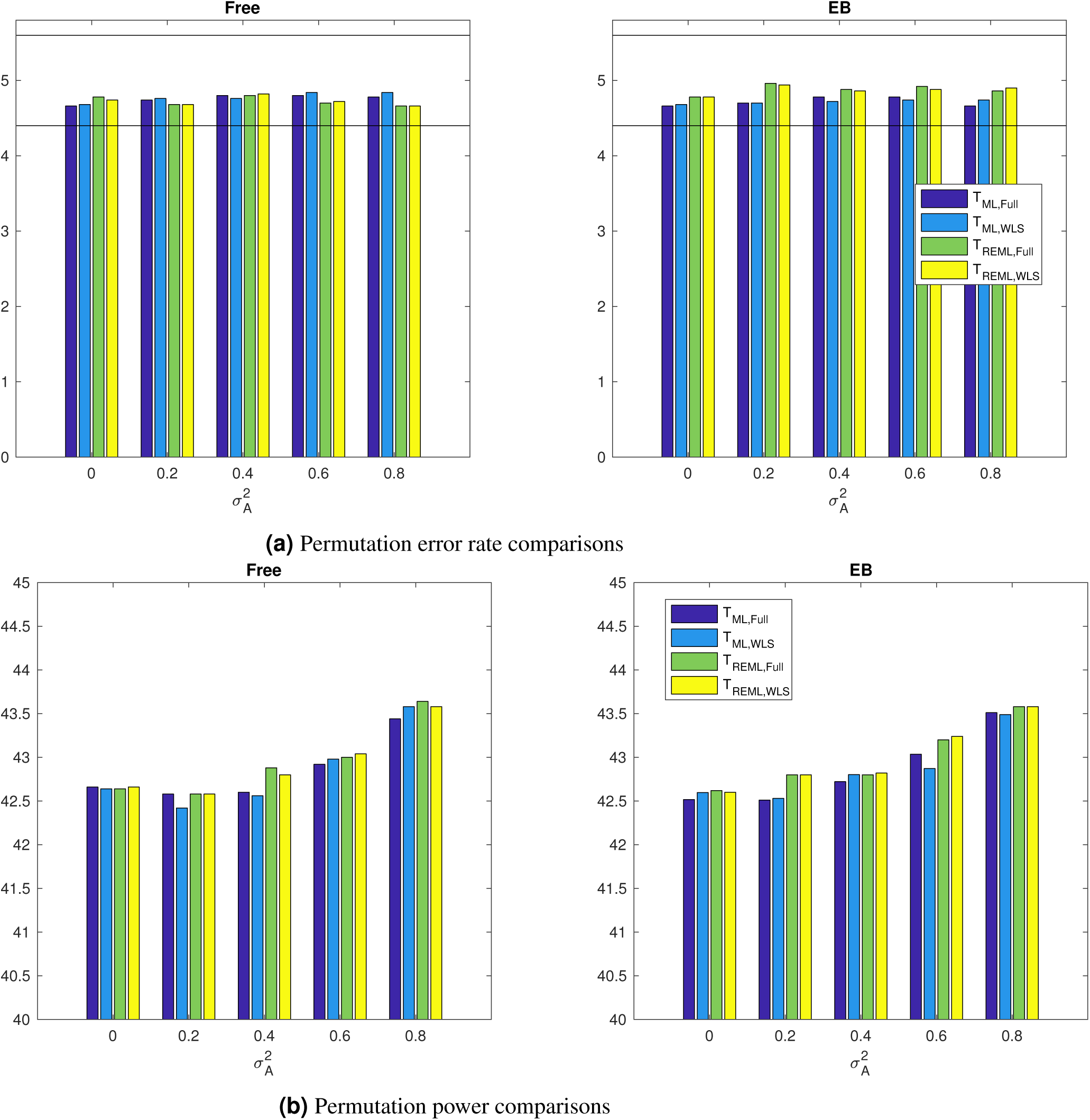
Simulation 2, comparing rejection rates of the proposed fixed effects permutation inference, for the null *β*_1_ = 0 (a) and alternative (b) for a 5% nominal level based on simulation using a GRM from 300 unrelated individuals and 5000 realizations and 500 permutations each realizations; left column shows results for the free permutation scheme, right for the exchangeability-block constrained method. Monte Carlo confidence interval is (4.40%, 5.60%). For non-iterative and fully converged, both permutation schemes could control the error rate at the nominal level, however free permutation is slightly more powerful than the constrained permutation.

Simulations show that the null distribution of the score test for *H*_0_: *β*_1_ = 0 based on the simplified models using the fully converged and non-iterative variance component estimators are valid and indistinguishable (Figure 2 & Supplementary Figure 6). However, we stress that the latter is much faster to calculate. Based on all of these results, we selected the score test based on the simplified REML function as the computationally most efficient test to be considered for genome-wide simulations and real data analysis.

**Figure 2.**
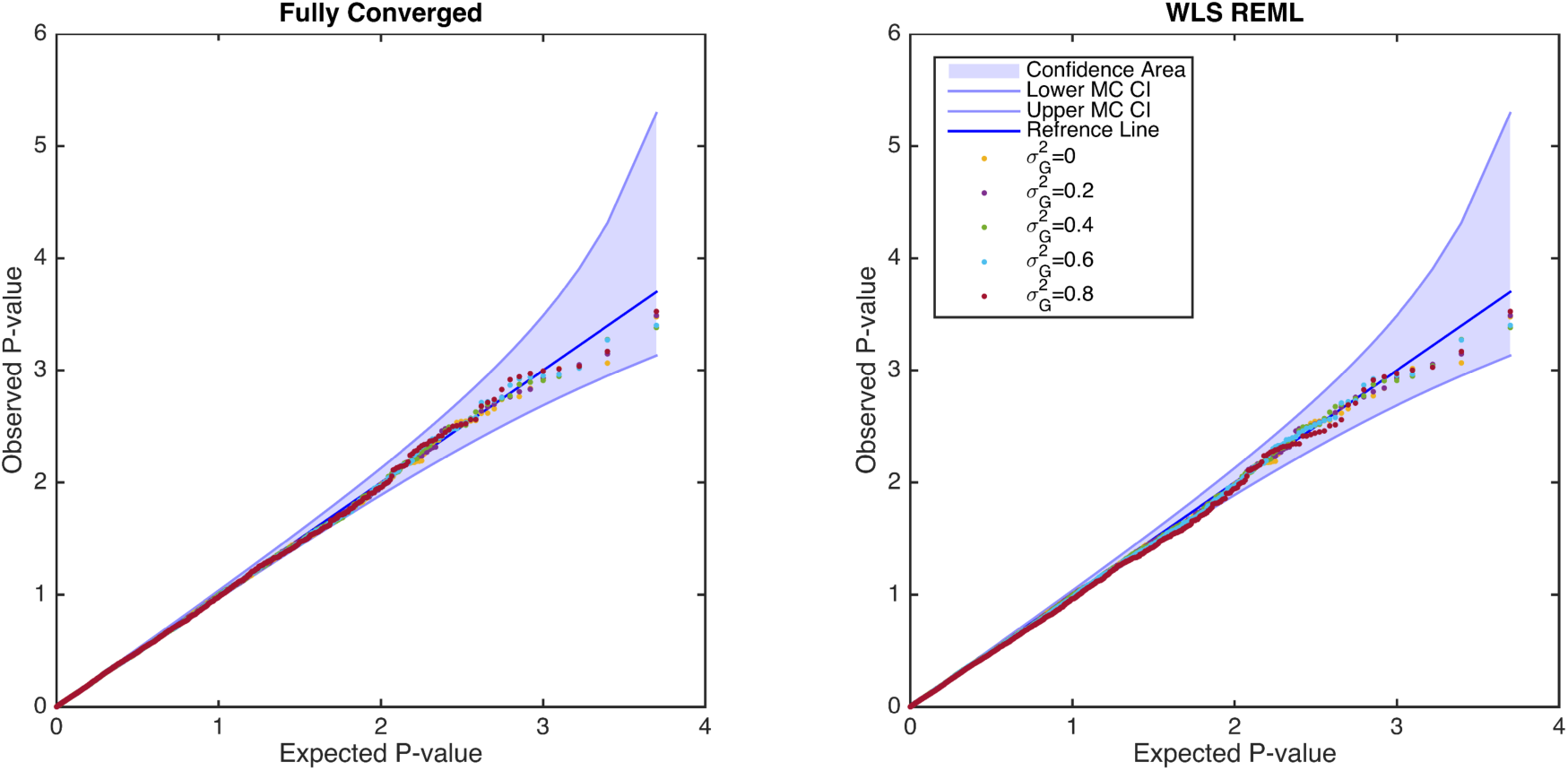
Simulation 3, comparing the distribution of null (*β*_1_ = 0) parametric p-values from the fixed effects score statistic derived from the simplified REML function using fully converged (left) and non-iterative (right) random effect estimator, for the GRM from 300 unrelated individuals. There is no apparent difference between the two random effect estimators, and both are consistent with a valid (uniform) P-value distribution. Confidence bounds created with the results of^41^ where ordered P-values follow beta distribution.

Genome wide simulations were conducted to compare the parametric P-values from FaST-LMM and the score test based on the simplified REML using non-iterative variance component estimator in terms of false positives and power. The simulation results reveal that both approaches provide overall valid error rates (FaST-LMM=4.94% and the WLS-REML score test = 4.89%, Figure 3a). Power simulation shows that FaST-LMM and the score test have largely similar power (FaST-LMM=15.25%, WLS-REML score test=15.22%), however, FaST-LMM is slower (Figure 3b). Despite reasonable concordance of P-value and fixed effect parameter estimates (*β*_1_) between FaST-LMM and simplified REML (Supplementary Figure 7), FaST-LMM’s estimates of parameter estimate variance 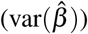 exhibits some systematic bias (Supplementary Figure 8).

**Figure 3.**
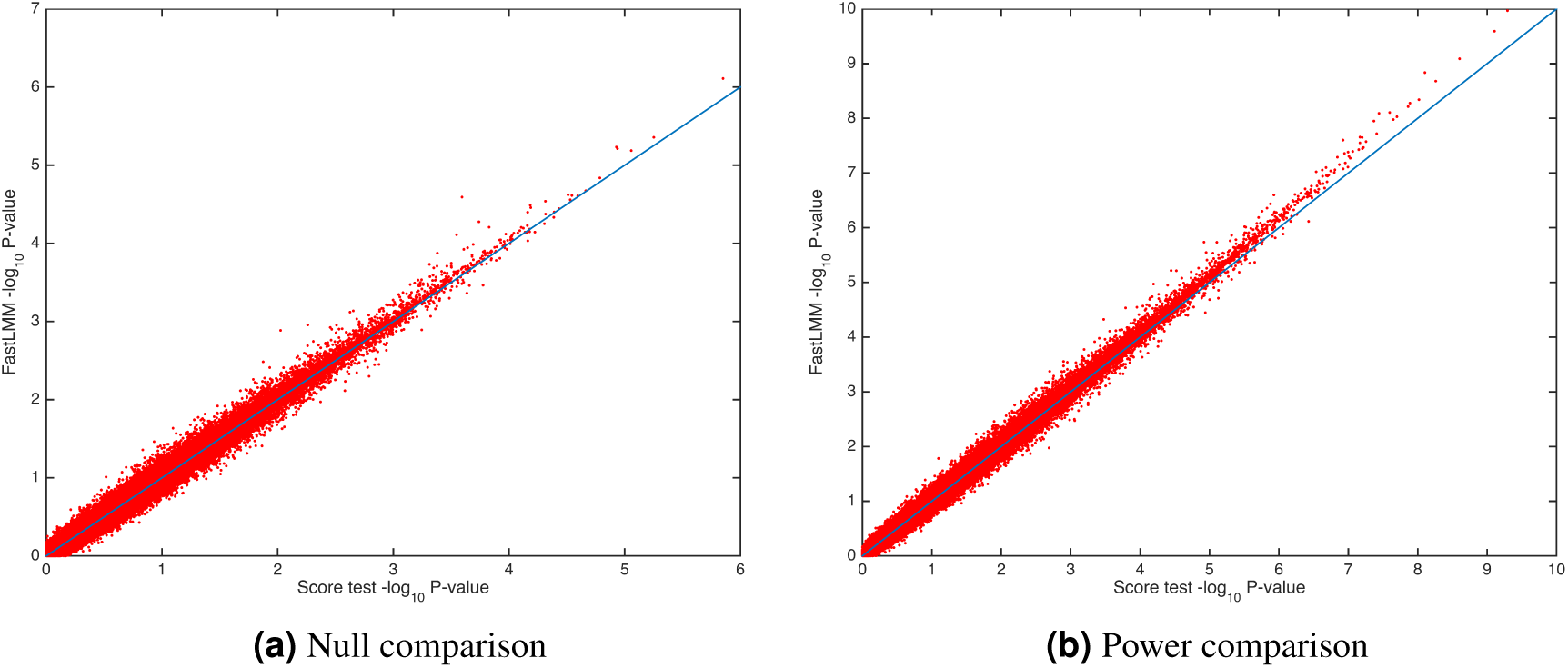
Simulation 4, comparing parametric p-values for simulated null data (a) and data with signal (b) using FaST-LMM’s LRT and our score test based on one-step optimization of the simplified REML function, using 100 random markers and 5000 realizations. The overall error rates for FaST-LMM and the score test are 4.94% and 4.89%, respectively, for nominal 5% where the Monte Carlo confidence interval is (4.40%,5.60%). While overall power is largely similar for both approaches (for FaST-LMM 15.25% and, and our method 15.22%), FaST-LMM is 200-fold slower.

**Figure 4.**
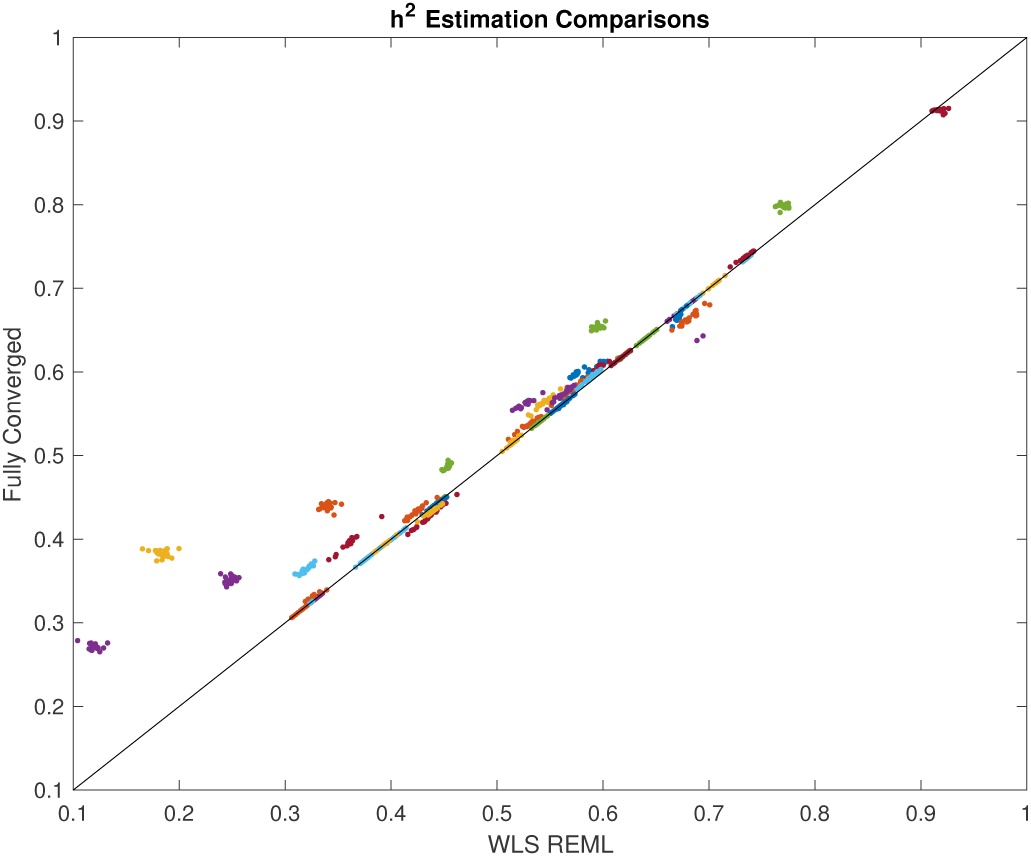
Real data analysis, comparing one-step and fully converged random effect estimators of 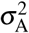 based on the simplified REML function. Colors represent random effect estimation at different regions for all 22 chromosomes. The scatter plot show consistent trend towards underestimation of random effect using non-iterative method, though this apparent increased accuracy comes with a 10^9^-fold greater computation time.

### Association Analysis of FA data

We performed GWA of whole brain fractional anisotropy (FA) data, using a whole brain parcellation of 42 regions of interest (ROIs) as well as a voxel-wise analysis for 53,458 voxels (332 subjects, 1,376,877 SNPs; for full details see Supplementary Methods), comparing the WLS-REML score test with the fully converged random effect estimators with FaST-LMM. We also evaluate the use of ordinary least square (OLS) with MDS as nuisance fixed effects regressors for control of population structure in GWA with unrelated individuals.

The random effect estimators, one-step and fully converged REML are compared directly in Figure (4) with a scatter plot, showing an apparent trade-off between accuracy and running time as the non-iterative method has lower estimates of 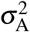 for some regions.

Even with the tendency for genetic variance to be underestimated with the non-iterative method, the association statistic show remarkable concordance, with both approaches having almost the same performance (Figure 5). FaST-LMM comparisons with the score test using the simplified REML function shows slightly larger statistics consistently for all ROIs, regardless of random effect estimation method (Supplementary Figures 10 & 11). Furthermore, comparing different approaches genomic control shows that regardless of random effect estimation method, the score test based on the simplified REML has smaller genomic control values than OLS with MDS nuisance regressors for all ROIs consistently. The genomic control of OLS with MDS nuisance regressors is poor, while the score test using both fully converged, one-step estimators and FaST-LMM have similar values close to unity (Supplementary Figure 9).

**Figure 5.**
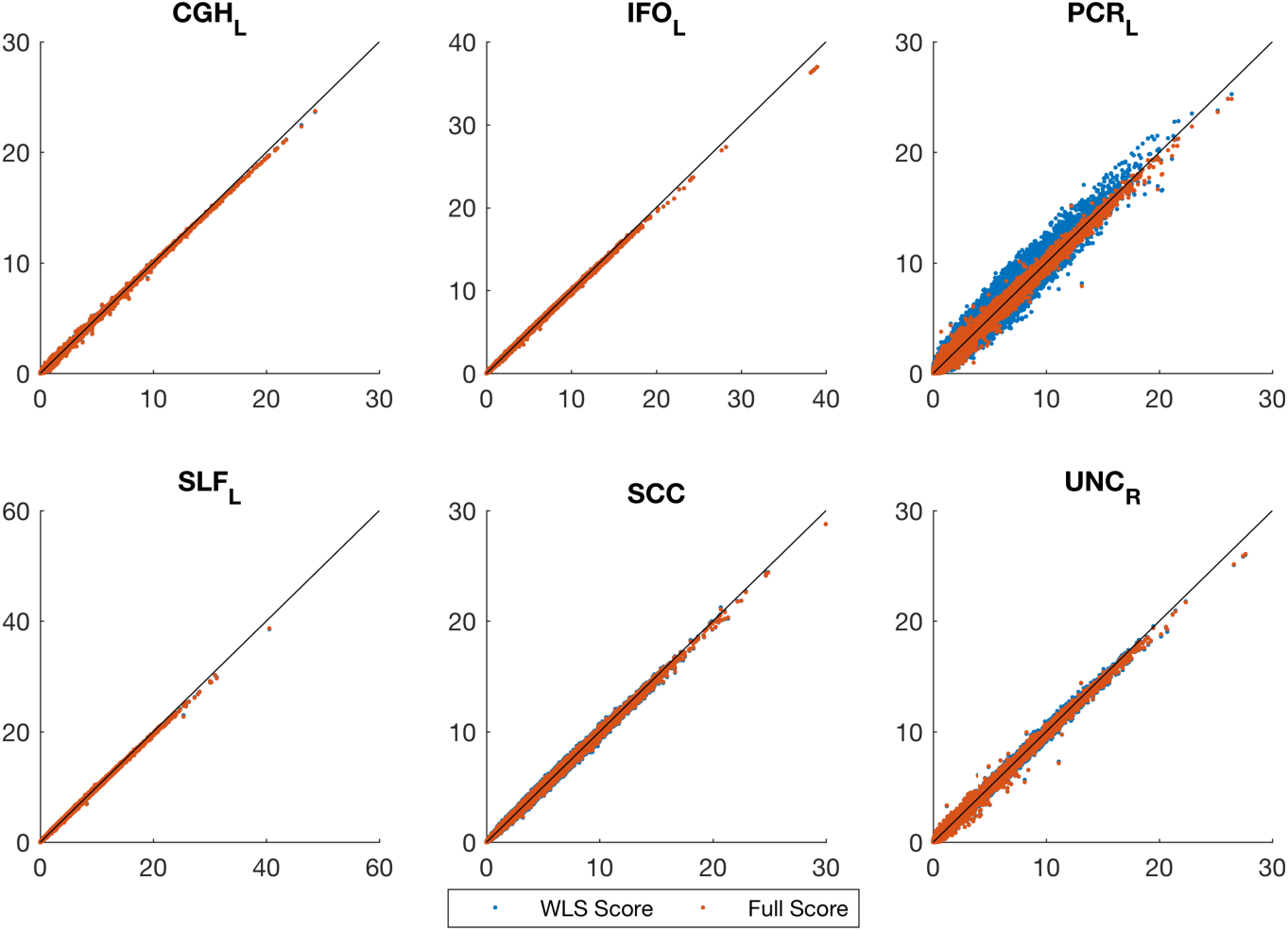
Real data analysis, comparing values of the score test for association testing (*H*_0_: *β*_1_ = 0) using non-iterative and fully converged random effect estimators and FaST-LMM’s LRT. Each plot represents a ROI where x-axis shows FaST-LMM’s LRT and y-axis represents the score test. Despite strong concordance between the score test results using WLS or fully converged random effect estimator, FaST-LMM is slightly more powerful, consistently for whole brain parcellation (see Supplementary Figures **??** & **??** for the rest of the ROIs).

Figure 6 compares QQ-plot of association statistics between our model, FaST-LMM and OLS with MDS. These plots show either an identical distribution or slightly larger values for the OLS approach; however, the OLS approach has poor genomic control (Supplementary Figure 9,) and after adjustment we get essentially identical results (Supplementary Figures 12 & 13).

**Figure 6.**
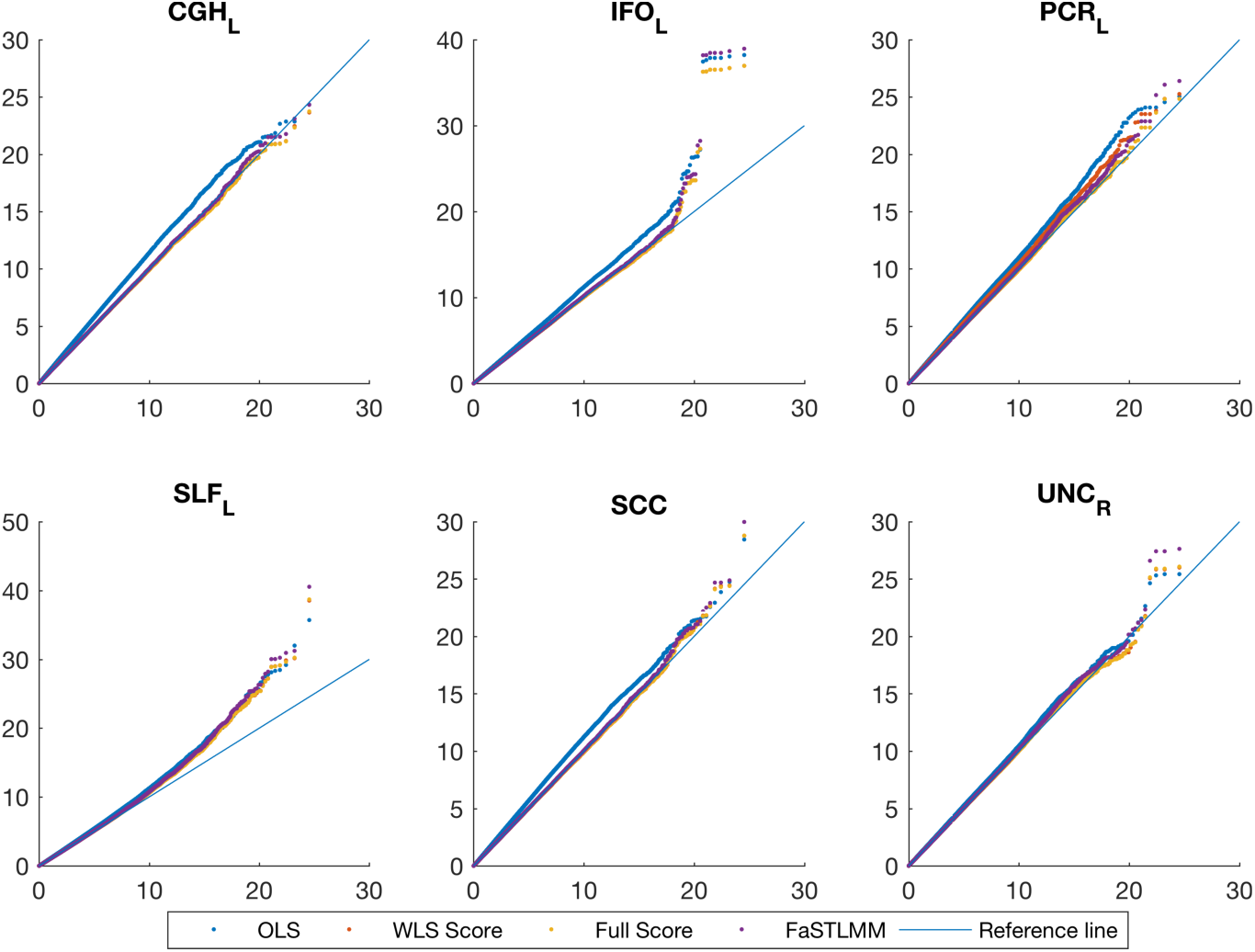
Real data analysis, QQ plot for comparing FaST-LMM and the score test based on the simplified REML function using the WLS-REML random effect estimator with the linear regression with MDS as nuisance fixed effects. Each plot corresponds to different ROIs. These plots show either an identical distribution or slightly larger values for the OLS approach. However the OLS approach has poor genomic control (Supplementary Figure 9).

A permutation test was used to find FWE-corrected P-values for 42 ROIs and 1,376,877 SNPs to assess association significance. Among the 42 *×* 1, 376, 877 *≈* 57 million statistics, 8 passed the permutation based FWE threshold 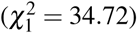. Application of a Bonferroni correction for 42 tests to the usual GWA alpha level (5 *×* 10^*-*8^) yields to a more stringent threshold 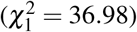 where only one association survives, indicating the potential improved power from a permutation-based inference that accounts for dependency among the tests (Figure 7).

**Figure 7.**
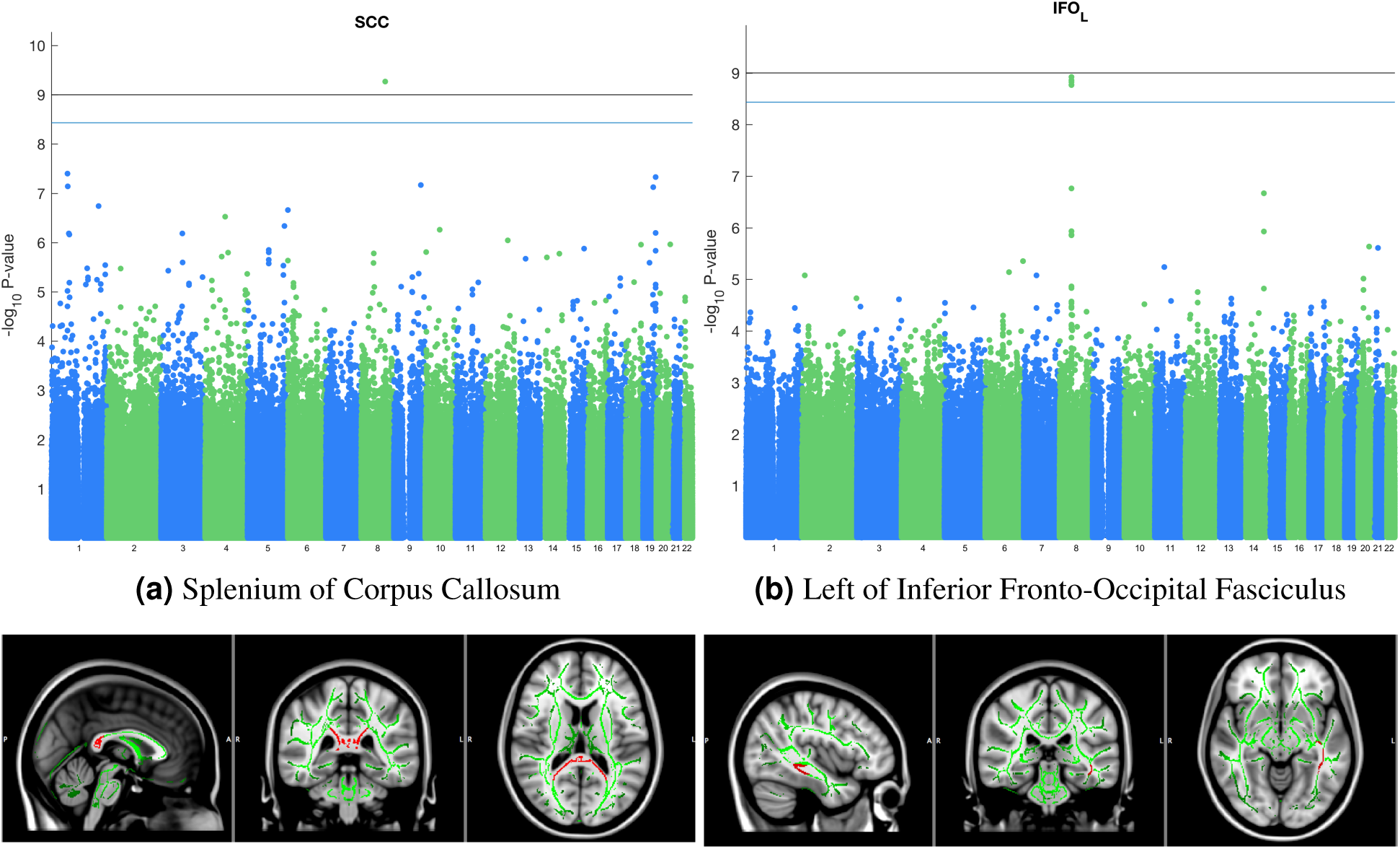
Real data analysis, GWA of whole brain fractional anisotropy data, using a whole brain parcellation of 42 regions. Permutation test was used to derive FWE corrected P-values of score test based on the simplified REML function using one-step random effect estimator. Among the 42 × 1, 376, 877 ≈ 57 million statistics, 8 passed the permutation based FWE threshold (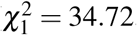, blue line in Manhattan plot). Application of a Bonferroni correction for 42 tests to the usual GWA alpha level (5 × 10^*-*8^) yields to a more stringent threshold (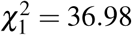, black line in Manhattan plot) where only one association survives, indicating the potential improved power from a permutation-based inference that accounts for dependency among the tests.

Finally we performed voxel-wise genome-wide association analysis of 53,458 voxels with 1,376,877 SNPs, using our proposed WLS-REML score test for association. Cluster-wise inference was performed on each spatial association map; we used a threshold corresponding to a 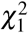 P-value of 0.01 to create clusters, and 1000 permutations were used to compute the maximum distribution of cluster size over space and SNPs, offering FWE control over the entire search space; voxel-wise FWE thresholds were also computed. The level 5% FWE-corrected voxel-wise statistic threshold was 66.42, producing 6 significant association out of 84 billion tests. The 5% FWE corrected cluster size threshold is 7370 but no SNP’s statistic map had a cluster exceeding this value; the largest observed cluster size is 6,648, which had a image-, genome-wide FWE-corrected cluster size P-value of 0.09.

### Benchmarking and running times

We compared running time of our WLS-REML score test to that of FaST-LMM, which to our knowledge is the fastest implementation of LMM. The comparison was done using simulated and read data with a Intel(R) core(TM) 3.4 GHz i7-2600 CPU and 16GB RAM. Parametric association testing of 5000 phenotypes with 6000 simulated markers using a sample of 300 individuals took 1 hour with FaST-LMM, however, our implementation of the score test (Eq. (4.12)) only took 3 seconds. On real data, parametric whole genome association on 22 ROIs, required 756 minutes using FaST-LMM while our approach took only 2 minutes.

## Discussion

Neuroimaging genetics has moved from establishing a heritable phenotypes to finding genetic markers that are associated with imaging phenotypes. Despite emerging world-wide consortia to boost GWA studies power using the largest possible sample sizes, there is a compelling need for powerful and computationally efficient analytic techniques that control for population structure at the site level.

Whether using the linear mixed model for controlling population structure or kinship, high dimensional imaging phenotypes presents challenges in terms of computational intensity and elevated false positive risk; growing sample sizes and whole genome sequence data add to the computational burden.

To tackle this problem, we used an orthogonal transformation that substantially reduced LMM complexity for GWA. The equivalence between projected model and REML function helped us to reduce complexity of association testing. Specifically, the projection reduces the information matrix to a scalar that enables efficient vectorized implementation of score test with time complexity *O*(*SNV*). Further improvements in speed can be achieved by using the WLS-REML random effect estimator with *O*(*NV*) that we found to be more accurate than the WLS-ML estimator.

We conducted intensive simulation studies, evaluating a broad set of test statistics for association testing using the simplified ML and REML functions accompanied by one-step and fully converged random effect estimators. The one-step random effect estimator using simplified REML function provides more accurate approximation of the fully converged one in comparison to the WLS-ML variance component estimator. The simulation and real data analysis shows that only minor differences in marker effect estimation and association test statistics between one-step and fully converged random effect estimator. However, the former requires less computational resources. Also, we could not observe any appreciable differences in performances in terms of the error rate and power using the GRM from unrelated individuals or kinship matrix from a family study.

The WLS-REML random effect estimator is fast enough to be used to estimate voxel-wise heritability. Although the proposed one-step random effect estimator is not as accurate as fully converged one, it can be used for filtering a small number of elements for further investigation with more computational intense tools. Furthermore, when restricted to individuals with European ancestry we found LMM had genomic controls values closer to 1 than OLS values, indicating the success of the LMM in dealing with population structure.

We selected the score test based on the simplified REML function for further investigations because it only requires a single variance component estimate, common to all markers under the null hypothesis. Furthermore, efficient vectorized implementation of score test for images accelerates association testing. The null distribution of WLS-REML score test P-values was nearly as accurate as for the fully converged REML score test, meaning that permutation is not required for element-wise inference.

Our contribution in the acceleration of the exact LMM can be seen at two steps. First, covariance matrix estimation using WLS-REML random effect estimator reduces time complexity from *O*(*N*^3^ + *INV*) to *O*(*N*^3^ + *NV*). Further improvement in speed is also obtained by using the score test based on simplified REML function. Our proposed method allows efficient implementation that reduces running time complexity to *O*(*SNV*). In addition, the efficient implementation of score test is fast enough to allow the permutation test to control family-wise error rate for number of elements in image and number of markers, and allow the use of spatial statistics like cluster size or TFCE.

Software implementation of this method, Nonparametric Inference for Genetics Analysis (NINGA), is available at http://nisox.org/software/ninga/.

## 1 Online Methods

At each voxel/element, a LMM for the genetic association for *N* individuals can be written as:

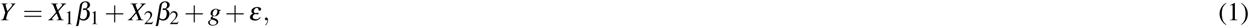

where *Y* is the *N*-vector of the measured phenotype; *X*_1_ is a *N*-vector of a given marker’s minor allele count, implementing an additive genetic model; *X*_2_ is the *N ×* (*P*1) matrix containing an intercept and fixed effect nuisance variables like age and sex; *β*_1_ is the scalar genetic effect; *β*_2_ is the (*P* 1)-vector of nuisance regression coefficients and *g* is the *N*-vector of latent (unobserved) additive genetic effects; and *ε* is the *N*-vector of residual errors. The trait covariance, var(*Y*)= var(*g* + *ε*)= ∑ can be written

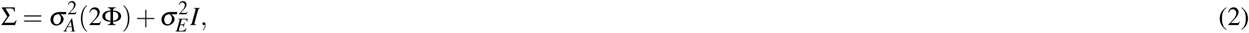

where 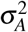 and 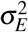 are the additive genetic and the environmental variance components, respectively; *I* is the identity matrix; and 2Φ is the GRM matrix where element (*i, j*) is calculated as:

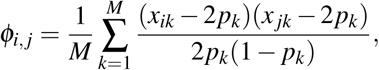

where *x*_*ik*_ is the minor allele count of the *i*-th subject’s *k*-th marker, coded as coded as 0, 1 or 2; *p*_*k*_ is frequency of the *k*-th marker; and *M* is the total number of markers.

Under the assumption that the the data follows a multivariate normal distribution, the model specified by Equations (2) and (1) have a log-likelihood of

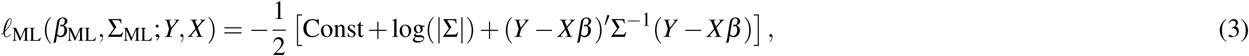

and a Restricted Maximum likelihood (REML) function of

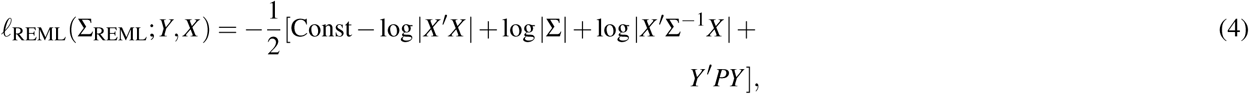

where *X* = [*X*_1_ *X*_2_] and *β* = [*β*_1_ *β*_2_] are the full design matrix of fixed effects and their parameter estimate vector, respectively, and *P* = ∑^*-*1^ (*I X* (*X ′* ∑^*-*1^ *X*)^*-*1^ *X ′* ∑^*-*1^), the projection matrix. The fixed effect parameters are estimated using generalized least squares (GLS)

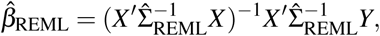

where 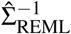 comes from optimised REML function (Eq. (4)).

Several algorithms have been proposed to accelerate ML or REML optimisation by transforming the model with the eigenvectors of the GRM and/or using a different covariance matrix parametrisation^21, 24, 33, 33 40^. Here we consider standard additive model covariance matrix parametrisation (Eq. (2)) as we can efficiently estimate it with our one-step, regression based approach^40^.

### 1.1 Simplified REML and ML Functions

The simplified ML function for LMM is discussed in^40^. For completeness, we review shortly the simplified ML function, to be next followed by development of the simplified REML function. The simplified ML function is obtained by transforming the data and model with an orthogonal transformation *S*, the matrix of eigenvectors of 2Φ that crucially coincide with the eigenvectors of ∑:

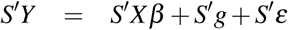

which we write as

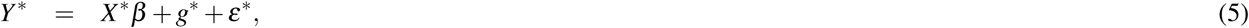

where *Y*^*∗*^ is the transformed data, *X*^*∗*^ is the transformed covariate matrix, *g*^*∗*^ and *ε*^*∗*^ are the transformed random components. The diagonalising property of the eigenvectors then gives a simplified form for the variance:

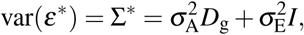

where ∑^*∗*^ is the variance of the transformed data and *D*_g_ = diag {λ_g*i*_} is a diagonal matrix of the eigenvalues of 2Φ.

The log likelihood takes on the exact same form as Equation (3) for *Y*^*∗*^, *X*^*∗*^, *β* and ∑^*∗*^, except is easier to work with since ∑^*∗*^ is diagonal:

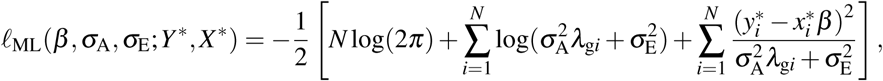

where 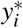 is the *i*-th element of *Y*^*∗*^, and 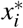 is the *i*-th row of *X*^*∗*^.

A simplified form of the REML function (Eq. (4)) is obtained by projecting the polygenic model (Eq. (1)) using eigenvector matrix of the adjusted GRM for the fixed effect nuisance terms as follows:

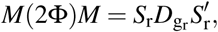

where *M* = *I X* (*X ′ X*)^*-*1^ *X′* is the residual forming matrix based on the fixed effects regressors; *D*_g_ r = diag *λ*_*gri*_ is the (*N − P*) (*N − P*) diagonal matrix of non-zero eigenvalues; and *S*_r_ is the *N ×* (*N − P*) matrix of eigenvectors that corresponds to non-zero eigenvalues. The projected polygenic model is obtained by pre-multiplying 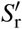 to the both sides of polygenic model (Eq. (1)):

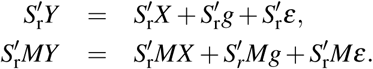

which we write as:

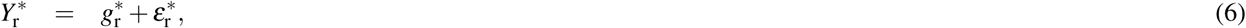

where 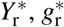 and 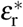 are *N − P* projected phenotype, genetic and residual vectors, respectively. In this fashion, the projected phenotype covariance matrix becomes diagonal:

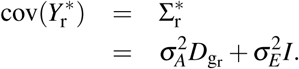

That is, the projected data, 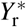, loglikelihood takes on a simpler form:

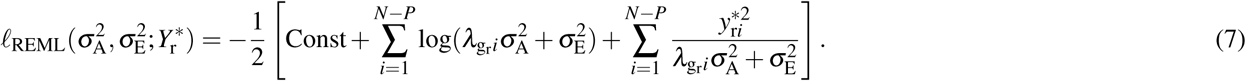

where 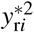 is the square of the *i*-th element of 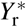.

In the **??** we show that this is equivalent to Equation (4), and thus we can use the eigenvectors of the adjusted GRM for nuisance fixed effects to build the REML log likelihood. It is clear from the Equation (7) that working with the simplified version of REML is computationally easier than the original one (Eq. (4)). Beside accelerating the REML optimisation, this approach facilitates performing likelihood ratio test for fixed effects (*β* s) and leads to a computationally efficient estimator and test statistic, described below.

### 1.2 REML and ML Parameter Estimation

We choose Fisher’s scoring method to optimize the simplified ML and REML functions because it leads to computationally efficient variance component estimators. The score and the expected Fisher information matrices for the simplified models can be expressed as:

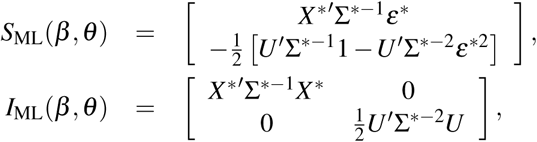

and

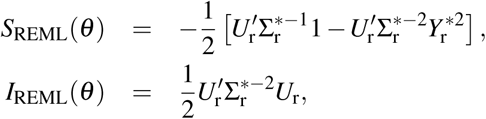

where 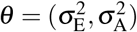; *U* = [**1**, *λ*_g_] and *U* = [**1**, *λ*_g_r__] are *N ×* 2 and (*N − P*) *×* 2 matrices; and *λ*_g_ is the vector of eigenvalues of (2Φ); *λ*_g_r__ the vector of eigenvalues of *M*(2Φ)*M*; **1** and **1**_r_ are *N* and (*N − P*)-vectors of one, respectively; 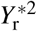 is the element wise product of 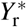; and *ε*^*∗*2^ is the element wise product of *ε*^*∗*^. Following Fisher’s scoring method it can be shown that at each iteration, maximum likelihood estimation of *β* and *θ* are updated based on WLS regression of *Y*^*∗*^ on *X*^*∗*^ and *ε*^*∗*2^ on *U*, respectively, as follows:

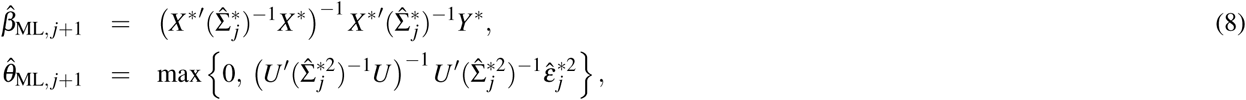

and restricted maximum likelihood estimation of *θ* is updated based on WLS regression of 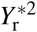 on *U*_r_ as follows:

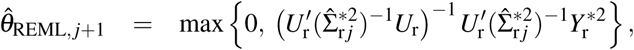

where *j* indexes iteration; 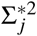 and 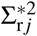 are constructed with *θ*_ML, *j*_ and *θ*_REML, *j*_ respectively; 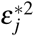 is the element-wise square of 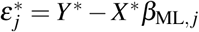; 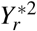 is the element-wise square of 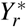; and the variance parameters *θ* must be positive, hence the maximum operator. As usual, these updates are iterated until convergence criteria holds.

It has been shown that when the initial value is a consistent estimator, the estimator based on the first iteration is asymptotically normal and consistent^42^. Such initial value for 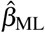 and 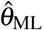 could be derived from OLS regression coefficients of *Y*^*∗*^ on *X*^*∗*^ and squared residuals on *U*, respectively:

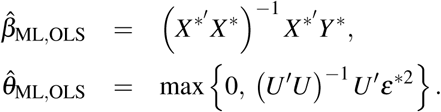

For REML, initial values for 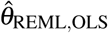 can be found as OLS regression coefficient of 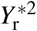 on *U*_r_:

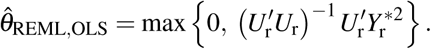

Hence our one-step, non-iterative estimators are:

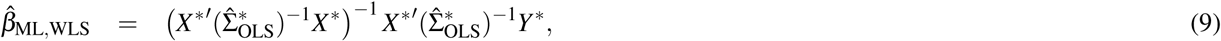

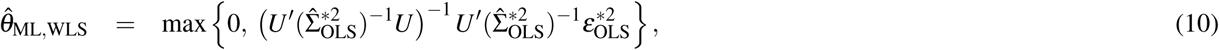

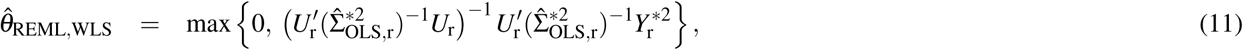

where 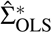 and 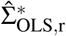 are formed by 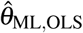 and 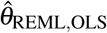 respectively, and 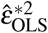 is the element-wise square of 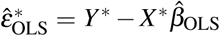. Going forward, we will use “ML” or “REML” to refer to the iterated estimators and “WLS” to refer to these one-step estimators.

### 1.3 Association Testing

The Score, likelihood ratio (LRT) and the Wald tests can be used for the genetic association testing using either ML or REML functions of the model in Equation (1).

The score statistic^43^ that requires the value of score and information matrices under the the null hypothesis constraint (*H*_0_: *β*_1_ = 0) for the simplified ML model (Eq. 5) can be written

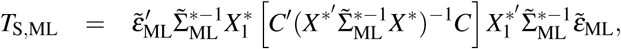

where *C* is a *P×* 1 contrast vector; 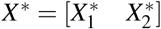 encompasses the full transformed covariate matrix; 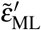 and 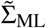 are the ML residual and covariance matrix estimates under the null hypothesis constraint. The score statistic for the projected model (Eq. (6)) can be derived like *T*_S,ML_ following the projection with respect to the *H*_0_ fixed effects, i.e. nuisance, terms *X*_2_,

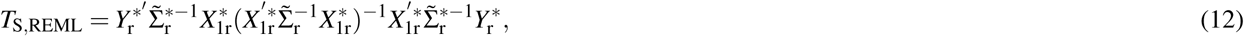

where 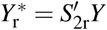 and 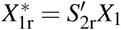 are (*N − P* + 1)-vectors of the projected phenotype and allele frequency count, respectively; and the projection matrix *S*_2r_ is comprised of the eigenvectors of *M*_2_ (2Φ)*M*_2_ with non-zero eigenvalues, 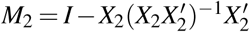; and 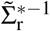 is the projected model covariance matrix estimation under the null model constraint.

**Likelihood Ratio Test** The LRT^44^ statistic is twice the difference of the optimised log-likelihoods, unrestricted minus *H*_0_-restricted. For ML this requires optimizing the likelihood function twice, once under the null *H*_0_: *β*_1_ = 0, once under the alternative. We denote the test statistic for this test *T*_L,ML_. A well-known aspect of REML is that it cannot be used to tests of fixed effects, since the null hypothesis would represent a change of the projection that defines the REML model. However, we can consistently use the same projection *S*_2r_, under the unrestricted and restricted models, to diagonalise our covariance and carry out such a hypothesis test. To be precise, the unrestricted model is

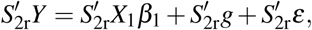

where 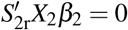 by the construction of *S*_2r_, and the restricted model is

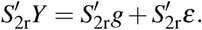

Following the same procedure as ML, the test statistic is denoted by *T*_L,REML_.

**The Wald Test** For a scalar parameter, the Wald test^43^ is the parameter estimate divided by the standard deviation of the estimate under an unrestricted model. For an vector parameter *β* and contrast *C*, it takes the form

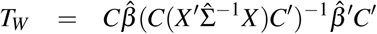

where 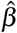 and 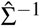 are the parameter estimations under the alternative hypothesis; this form holds for both ML and REML. A test for genetic association testing can be calculated either using fully converged or one-step variance component estimators. In the parametric framework, all of the aforementioned tests null distribution follow chi square distribution with one degree of freedom asymptotically.

### 1.4 Inference Using Permutation Test

In neuroimaging the permutation test is a standard tool to conduct inference while controlling the family wise error rate (FWE)^39^. It only requires an assumption of exchangeability, that the joint distribution of the error is invariant to permutation, and can be provide exact inference in the absences of nuisance variables, or approximately exact control with nuisance variables^45^. Control of the FWE of a voxel-wise or cluster-wise statistic is obtained from a maximum distribution of the corresponding statistic. However naive use of permutation test for the genetic association testing, ignoring dependence structure between individuals, leads to invalid inferences^46, 47^. Here we propose two permutation schemes that account for dependence explained by our model, one free and one constrained permutation approach.

#### Free Permutation

The genetic association testing in the context of LMM using a permutation test requires proper handling of fixed effect and random effect nuisance variables in order to respect the exchangeablity assumption. While there are a variety of methods for testing for a fixed effect when the errors are independent^45^. However little work has been done for fixed effect inference using permutation test in linear mixed models where the error term is correlated^46^.

**Free Permutation for the simplified ML Model:** For the simplified model (Eq. 5) we create permuted data 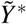 using the reduced, *H*_0_: *β*_1_ = 0 null model residuals and use them to create surrogate null data,

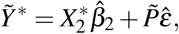

where 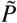 is one of *N*! possible *N × N* permutation matrices; 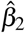 is the reduced model nuisance estimate found with either fully converged (Eq. (8)) or one-step (Eq (9)) methods; 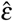 denotes the reduced model residuals likewise found with either fully converged or one-step estimators; and the tilde accent on the data 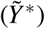 and permutation 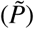 denotes one of many null hypothesis realisations. The reduced null model is not exchangeable due heteroscedasticity of ∑^*∗*^, but we account for this in each permutation step by fitting the simplified model (5) with the permuted covariance matrix

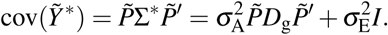

With this approach we obtain samples from the empirical null distribution of the maximum score, LRT and the Wald tests (or cluster-size, after thresholding one of these test statistics), where the maximum is taken over all voxels and SNPs to control FWE.

**Free Permutation for Simplified REML model**: While above, we created null hypothesis realizations by permuting the nullmodel residuals and adding back on estimated nuisance, here we will permute data after reduced-model eigen-transformation. We do this because projection removes the nuisance fixed effect covariates. In both cases, though, we must account for the dependence existing under *H*_0_.

An alternate permutation scheme could be built based on projecting the LMM model (Eq. (1)) to the lower dimension space with respect to the null hypothesis reduced model, i.e. using only the nuisance fixed effect terms. Let *M*_2_ be the residual forming based on *X*_2_ alone and, as Section 1.1, define *S*_2r_ as the transformation based on the non-trivial eigenvectors of *M*_2_ (2Φ)*M*_2_, creating a model with dimension N-(P-1):

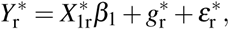

where 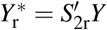 is the reduced transformed data; 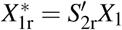 is as above, the reduced transformed additive genetic effect; 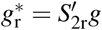 and 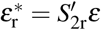 are the latent genetic effect and random error terms, respectively, after the reduced transformation. Here we permute the data, producing 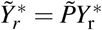, with permuted covariance matrix

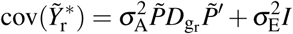

fit in each permutation step. However, under the null hypothesis of no genetic effect, estimated random effects for permuted phenotype are exactly as same as the un-permuted phenotype and hence the variance components only need to be estimated once.

#### Constrained Permutation: exchangeability blocks

In the free permutation approaches we permute despite the lack of exchangeability, but then permute the covariance structure to account for this. An alternate approach is to define exchangeability blocks where observations within each block can be regarded as exchangeable. Precisely, with exchangeability blocks, the assumption is that permutations altering the order of observations only within each block preserve the null hypothesis distribution of the full data.

While not feasible for the original correlated model (1), in the simplified ML (5) or simplified REML (6) model we can define approximate exchangeability blocks. In simplified models the sorted eigenvalues arrange the observations by variance (increasing or decreasing, depending on software conventions). Hence blocks of contiguous observations *Y*^*∗*^ or 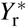 will have variance that is as similar as possible and will be, under the null hypothesis, approximately exchangeable.

We propose to define exchangeability blocks such that the range of *D*_g_ or *D*_g_ r values within a block is no greater than 0.01. This cut off ensured the eigenvalues did not vary by more than a factor of 10% within a block. Permutation is constrained within these blocks and the test procedure is as described above for simplified ML and REML free permutation schemes above, except that the test statistic is computed using the unpermuted covariance matrix.

### 1.5 Efficient score statistic implementation for vectorized images

To fully exploit the computational advantage of our non-iterative, reduced-dimension projected model estimation method we require a vectorized algorithm. That is, even without iteration, the method will be relatively slow if the evaluation of the estimates is so complex that each phenotype must be looped over one-by-one. For fast evaluation with a high-level language like Matlab, the estimation process for a set of phenotypes must be cast as a series of matrix algebra manipulations.

In this section we develop the vectorized algorithm for association one chromosome’s worth of SNPs and all image voxels/elements (subject to memory constraints). To avoid proximal contamination^24^ and efficient implementation of LMM, we follow leave one chromosome out approach where all markers on a chromosome being tested are excluded from the GRM^30, 32^.

Let **Y**_r_ and **X**_r_ be a (*N − P*) × *V* and (*N − P*) × *G* matrices of projected traits and allele frequencies respectively, where *V* and *G* are number of elements in image and number of SNPs the tested chromosome, respectively. The score test requires parameter estimation under the null hypothesis constraints, and since *X*_2_ is the same for all SNPs, the estimated covariance matrix will be the same all markers the chromosome. Thus the covariance matrix only need to be estimated once as follows:

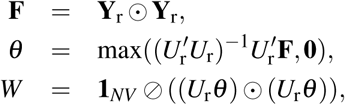

where **F** and **Y**_r_ are (*N − P*) *×V* matrices, where each column of **Y**_r_ is 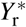 (Eq. (6)) for one image element and **F** is the element-wise squaring of **Y**_r_; ⊙ denotes Hadamard product; ⊘ denotes element-wise division; *θ* is the 2 *×V* matrix of OLS solutions which is matrix counterpart of 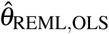; **0** is the 2 *×V* matrix of zeros; and here max(*·,·*) is an element-wise maximum between the two operands, evaluating to a 2 *×V* matrix; *W* and **1**_*NV*_ are the (*N − P*) *×V* matrices, where each column of *W* is 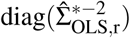 for the corresponding image element and **1**_*NV*_ is a matrix of ones. With the following notation,

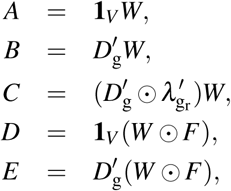

where **1**_*V*_ is the length-*V* column vector of ones, we can compute the variance components of the vectorized image as:

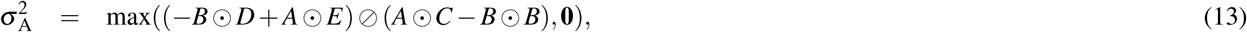

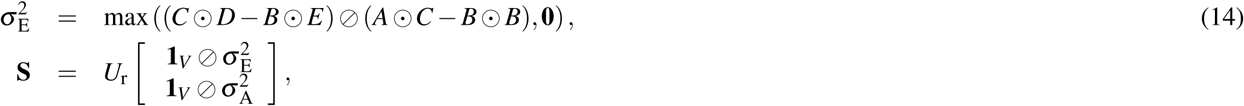

where 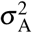 and 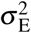 are the length-*V* column vectors of genetic and environmental variance components, respectively; and **S** is a (*N-P*) *× V* matrix which here each column of **S** is the element-wise reciprocal of the diagonal of the variance matrix of the corresponding image element’s data **Y**_r_ for each element of image. In this fashion, the score statistic matrix for all markers being tested and the vectorized image can be expressed as:

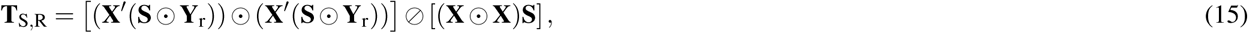

where **T**_S,R_ is a *G×V* matrix of score statistics for all SNPs and traits.

## Acknowledgements

TEN is supported by the Wellcome Trust. HG, TEN & PK were supported by National Institutes of Health R01 EB015611-01.

## Author contributions statement

H.G. developed the statistical method, ran the simulations and analyzed the simulated and real data. P.K. provided the human data and supervised its analysis. B.D. implemented the method both in CPU and GPU. A.M.W., D.C.G. and J.B helped in writing the paper. T.E.N. supervised the project. All authors contributed to critical review of the manuscript during its preparation.

## 1 Supplementary Methods

### 1.1 Simplified REML Function

At each voxel/element, a LMM for the genetic association for *N* individuals can be written as:

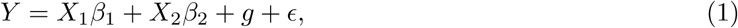

where *Y* is the *N*-vector of the measured phenotype; *X*_1_ is a *N*-vector of a given marker’s minor allele count, implementing an additive genetic model; *X*_2_ is the *N×* (*P* − 1) matrix containing an intercept and fixed effect nuisance variables like age and sex; *β*_1_ is the scalar genetic effect; *β*_2_ is the (*P* 1)-vector of nuisance regression coefficients and *g* is the *N*-vector of latent (unobserved) additive genetic effects; and *ϵ* is the *N*-vector of residual errors. The trait covariance, var(*Y*) = var(*g* + *ϵ*) = ∑ can be written

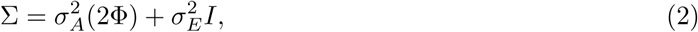

where 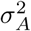 and 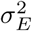 are the additive genetic and the environmental variance components, respectively; *I* is the identity matrix; and 2< I is the GRM matrix.

Under the assumption that the the data follows a multivariate normal distribution, the model specified by Equations (2) and (1) have a Restricted Maximum likelihood (REML) function of

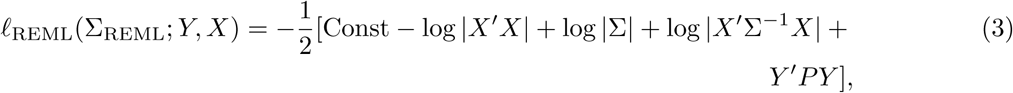

where *X* = [*X*_1_ *X*_2_] and *β* = [*β*_1_ *β*_2_] are the full design matrix of fixed effects and their parameter estimate vector, respectively, and *P* = ∑^*-*1^ (*I X*(*X′* ∑^*-*1^ *X*)^*-*1^ *X′* ∑^*-*1^), the projection matrix. The fixed effect parameters are estimated using generalized least squares (GLS)

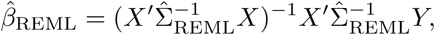

where 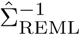 comes from optimised REML function (Eq. (3)).

It is clear from Equation (3) that computational burden of REML function is substantial even for small datasets. Here we introduce an orthogonal transformation that accelerates the REML function optimisation dramatically. To obtain such transformation, we first show that the likelihood function of transformed data with any orthogonal residualising matrix, that is a matrix that maps *Y* to the null space of *X*, is exactly the same as the REML function. Then, a simplifed form of the REML function (Eq. (3)) can be obtained using a particular transformation that makes the covariance matrix of transformed data diagonal.

Let *M* = *I X*(*X′ X*)^*-*1^ *X′* be the residual forming matrix based on the fixed effects regressors. Since *M* is idempotent, it can be diagonalised

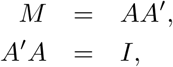

where *A* is the *N ×* (*N − P*) matrix of the eigenvectors of *M* corresponding to the non-zero eigenvalues. Crucially, *A* also residualises the data, because it is orthogonal to the design matrix *X*:

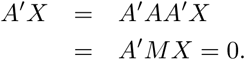

Hence *A′ Y ∼ N* (0, *A′* ∑*A*) and the log likelihood of the transformed data is

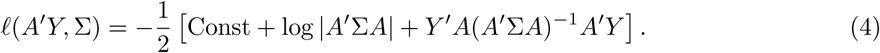

Now we show that this (Eq. (4)) is equivalent to Equation (3), and thus we can use the eigenvectors of the residual-forming matrix to build the REML log likelihood. Assuming that *X* is a full column rank, then [*A X*] is the *N × N* square non-singular matrix. To show that the project data loglikelihood is the same as the REML function (Eq. (3)) we use the following identity:

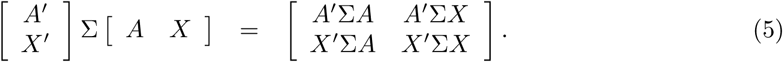

Taking the determinant of LHS of the Equation (5) gives us:

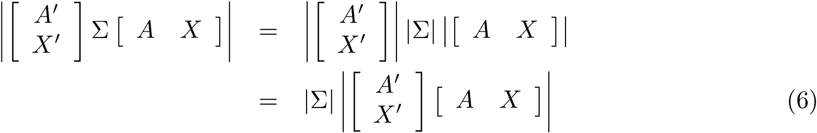

The RHS determinant of the Equation (5) using the block matrix determinant rule can be written as

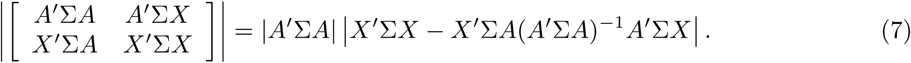

Hence taking the determinant from the identity in the Equation (5) gives us

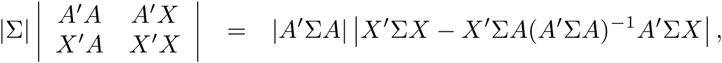

where using *A′ A* = *I*, *A′ X* = 0, it can be shown that

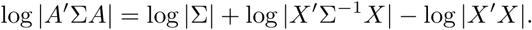

Finally, using *A*(*A′* ∑*A*)^*-*1^ *A* = *P* [Searle et al., 2009, M4.f p. 451], it is clear that Equations (3) and (4) are equivalent. Hence, transformed data likelihood function is exactly as same as the REML function.

As *A* is not unique, we seek to find one that diagonalises the covariance of the residualised data. The transformation matrix could be derived from eigendecomposition of GRM adjusted for the fixed effect covariates as follows:

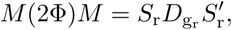

where 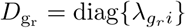 is the (*N − P*) *×* (*N − P*) diagonal matrix of non-zero eigenvalues; and *S*_r_ is the *N ×* (*N − P*) matrix of eigenvectors that corresponds to non-zero eigenvalues. Firstly, *S*_*r*_ is a valid *A*, because its columns are orthogonal 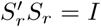 and 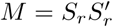 [Kang et al., 2008]. Thus we define the projected polygenic model by pre-multiplying 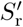 both sides of polygenic model (Eq. (1)):

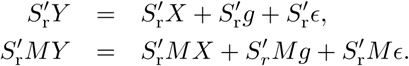

which we write as:

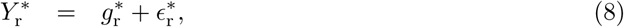

where 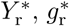 and 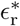 are *N − P* projected phenotype, genetic and residual vectors, respectively. In this fashion, the projected phenotype covariance matrix becomes diagonal:

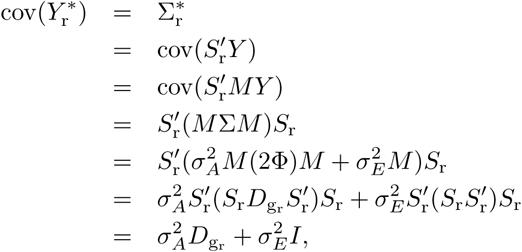

where we have used the identity 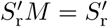. That is, therefore the projected data, 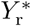, loglikelihood takes on a simpler form:

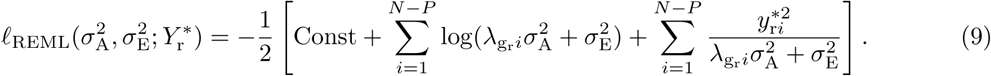

where 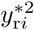 is the square of the *i*-th element of 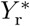.

### 1.2 Simulations

Intensive simulation studies are conducted to evaluate proposed methods for association estimation and testing. The aim of the first study is to compare fully converged and one-step random effect estimators based on the simplified ML and REML functions. In the second study, the performance of various test statistics for the association testing are compared using a fully converged or one-step random effect estimators for ML and REML functions. Finally, we compare FaST-LMM [Lippert et al., 2011] to our preferred test, the score test based on the simplified REML function, **T**_S,REML_, using both false positive error rates and empirical power using simulated genetic markers.

In all simulations the response variable is assumed to follow *Y* = *Xβ* + *ϵ*, where *ϵ ∼ N* (0, ∑) and 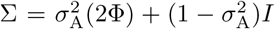, giving a unit variance phenotype. As above, the design matrix is partitioned *X* = [*X*_1_ *X*_2_], where *X*_1_ is the allele count per subject for a given marker, and *X*_2_ are all other non-genetic fixed effects. In our simulations, *X*_1_ is based on simulated marker, where each marker has a reference allele frequency sampled from a uniform distribution on [0.1, 0.9]. The *X*_2_ matrix has 3 columns, an intercept, a linear trend from −1 to 1, and the element-wise square of the linear trend. Kinship matrices from a family study, genetic analysis workshop 10 (GAW10), and genetic similarity matrix from simulated genetic markers for a sample of unrelated individuals with different sizes were chosen to set the covariance, for a range of genetic variances, 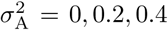. Specifically, the Cholesky decomposition of ∑ was used to premultiply i.i.d normal random variables with 5000 realisations.

### 1.3 Genome wide simulations

FaST-LMM and the score test performances based on P-value, parameter estimate 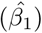 and variance of parameter estimate 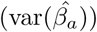 are compared using simulated SNPs and phenotype when there is no population structure. 60,000 SNPs for 300 individuals with minor allele frequency between (0.05, 0.5) were simulated. 6000 null and 100 causal markers were used to compare the false positive rates in 5000 realisations. In the null simulations 6000 markers were used to induce different level of heritability under the additive model 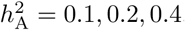 and 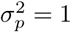 where markers are standardised to have mean zero and unit variance. Finally for power simulations 100 markers were explained 30% of phenotipic variance where the effect for each marker was drawn from *N* (0, 0.3*/*100).

### 1.4 Real Data

To validate our proposed methods for association estimation and inference for imaging data, we applied them on a dataset from healthy and schizophrenic individuals to perform ROI and voxelwise genome wide association analysis using cluster wise inference. The sample was 54% healthy individual (175 control/155 schizophrenic) and had a mean age of 39.12 (SD= 14.9) where 50% of the sample is male.

#### Diffusion Tensor Imaging

Imaging data was collected using a Siemens 3T Allegra MRI (Erlangen, Germany) using a spinecho, EPI sequence with a spatial resolution of 1.7 *×* 1.7 *×* 4.0 mm. The sequence parameters were: TE/TR=87/5000ms, FOV=200mm, axial slice orientation with 35 slices and no gaps, twelve isotropically distributed diffusion weighted directions, two diffusion weighting values (b=0 and 1000 s/mm2), the entire protocol repeated three times.

ENIGMA-DTI protocols for extraction of tract-wise average FA values were used. These protocols are detailed elsewhere [Jahanshad et al., 2013] and are available online http://enigma.ini.usc.edu/protocols/dti-protocols/. Briefly, FA images from subjects were non-linearly registered to the ENIGMADTI target brain using FSL’s FNIRT [Jahanshad et al., 2013]. This target was created as a minimal de-formation target based on images from the participating studies as previously described (Jahanshad et al., 2013b). The data were then processed using FSL’s tract-based spatial statistics (TBSS; http://fsl.fmrib.ox.ac.uk/fsl/fslwiki/TBSS) analytic method [Smith et al., 2006] modified to project individual FA values on the hand-segmented ENIGMADTI skeleton mask. The protocol, target brain, ENIGMADTI skeleton mask, source code and executables, are all publicly available (http://enigma.ini.usc.edu/ongoing/dti-working-group/). The FA values are normalized across individuals by inverse Gaussian transform [Servin and Stephens, 2007, Allison et al., 1999] to ensure normality assumption. Finally, we analyzed the voxel and cluster-wise FA values with applying along the ENIGMA skeleton mask.

#### Genetic Quality Control

In this study only genotyped Single Nucleotide Polymorphisms (SNPs) from genome-wide information were included in the analysis. Visual inspection of multi-dimensional scaling analysis was used to extract individuals with European ancestry. SNPs from individuals with European ancestry that do not meet any of the following quality criteria were excluded: genotype call rate 95%, significant deviation from HardyWeinberg equilibrium *p<* 10^*-*6^ and minor allele frequency 0.1 was used to ensure that sufficient numbers of subjects would be found in our sample in each genotypic group (homozygous major allele, heterozygous, homozygous minor allele) using an additive genetic model. After all quality control steps, 962,885 out of 1000,000 SNPs remain for genome-wide association analysis.

## 2 Supplementary Figures

**Figure 1:**
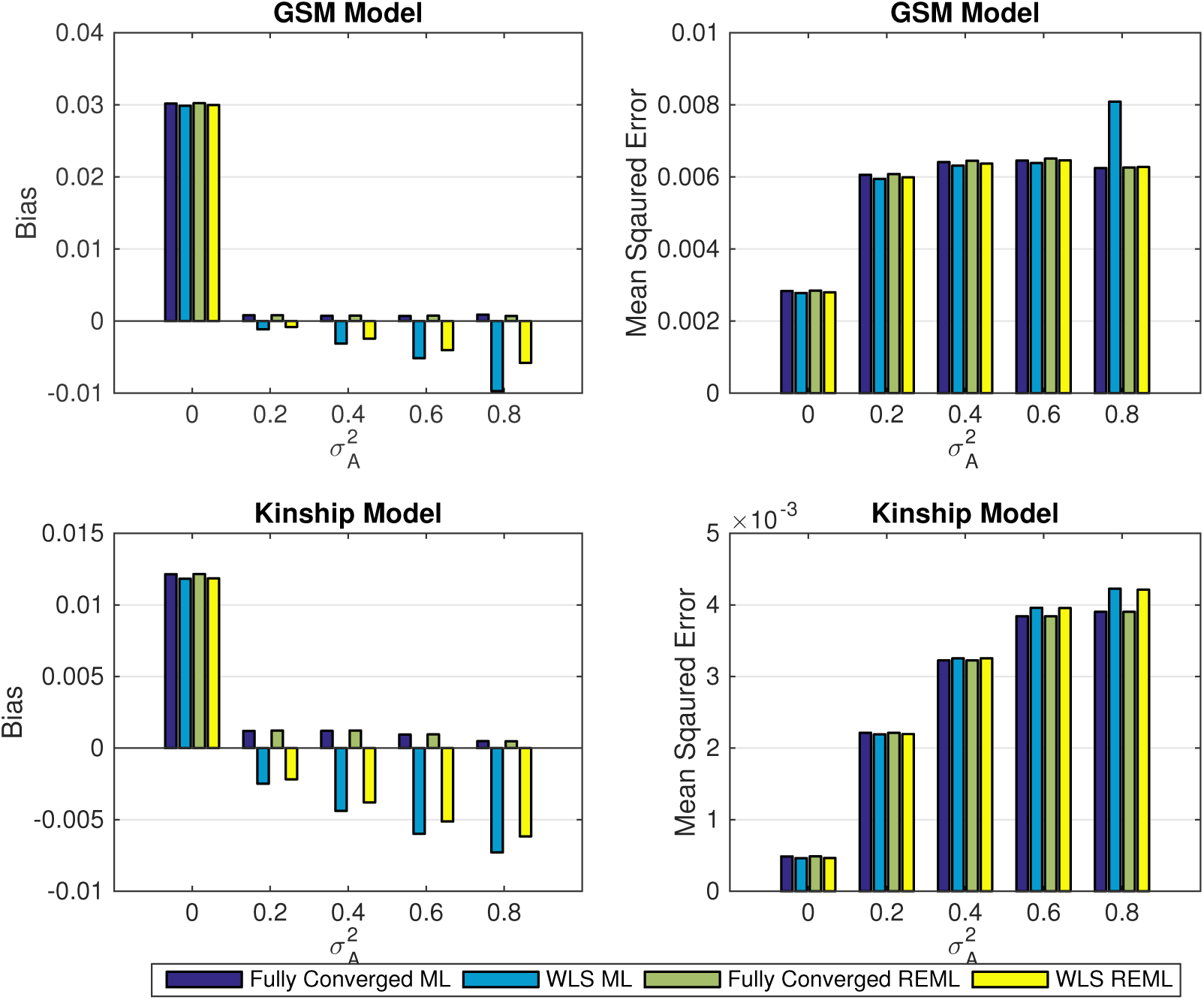
Simulation 1 results, comparing the bias (left column) and mean squared error (right column) of non-iterative and fully converged random effect estimators using the simplified ML or REML for 5000 realizations, for different level of genetic random effect 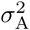. The results are based on a GRM constructed from 1200 unrelated individuals (top row) and kinship matrix from GAW 10 with 23 families and 1497 individuals (bottom row). While the one-step estimators generally (ML or REML) have more bias than fully converged ones, WLS-REML has less bias than WLS-ML, and in terms of MSE there is a relatively small difference in performance among all the methods.

**Figure 2:**
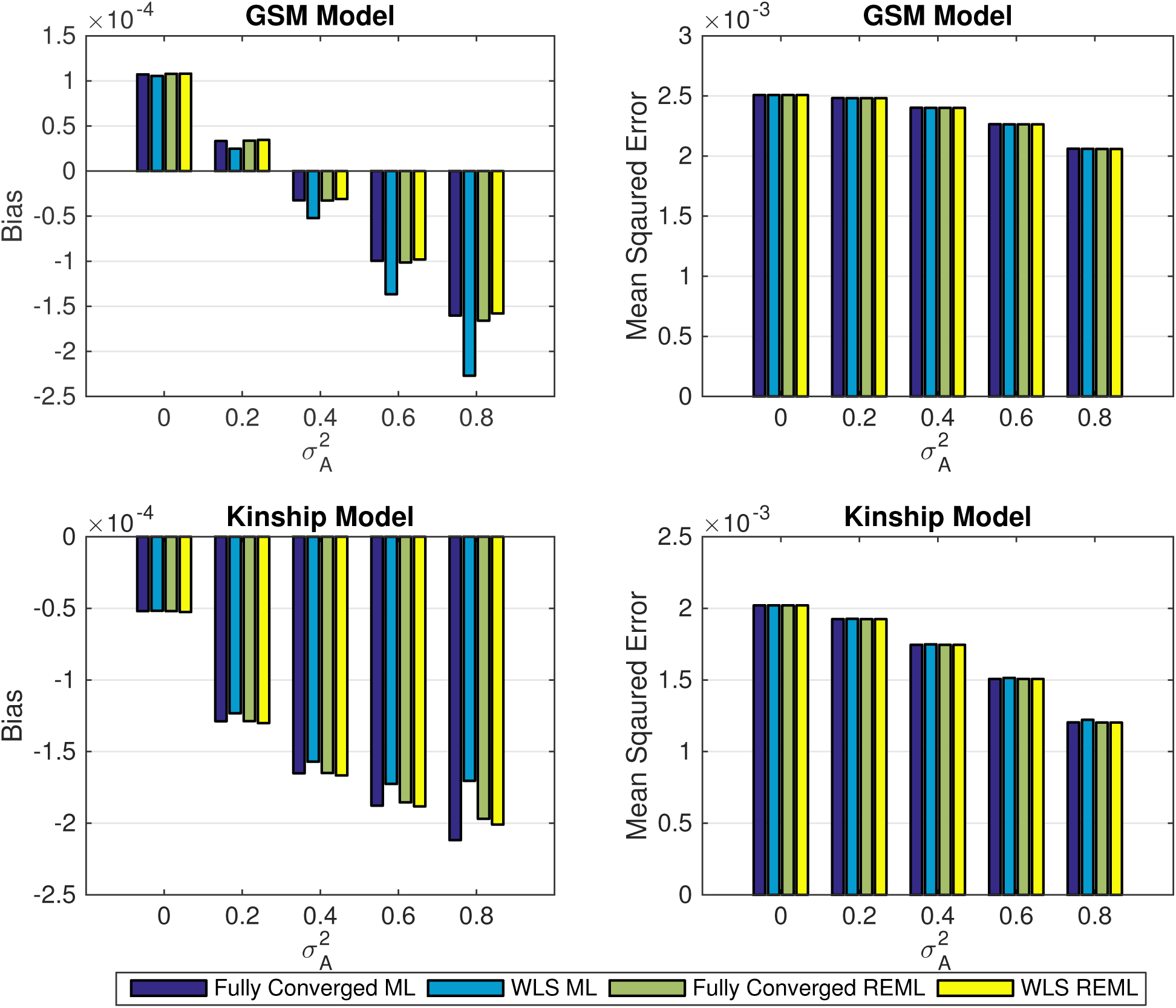
Simulation 1, comparing the bias (left column) and mean squared error (right column) of the fixed effect (additive allelic effect, *β*_1_) using the simplified ML or REML for 5000 realizations, for different level of genetic random effect 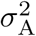 when *β*_1_ = 0. The results are based on a GRM constructed from 1200 unrelated individuals (top row) and kinship matrix from GAW 10 with 23 families and 1497 individuals (bottom row). These results show that fixed effect estimation using WLS-REML variance component estimator has almost identical performance as the fully converged one over different levels of genetic variance.

**Figure 3:**
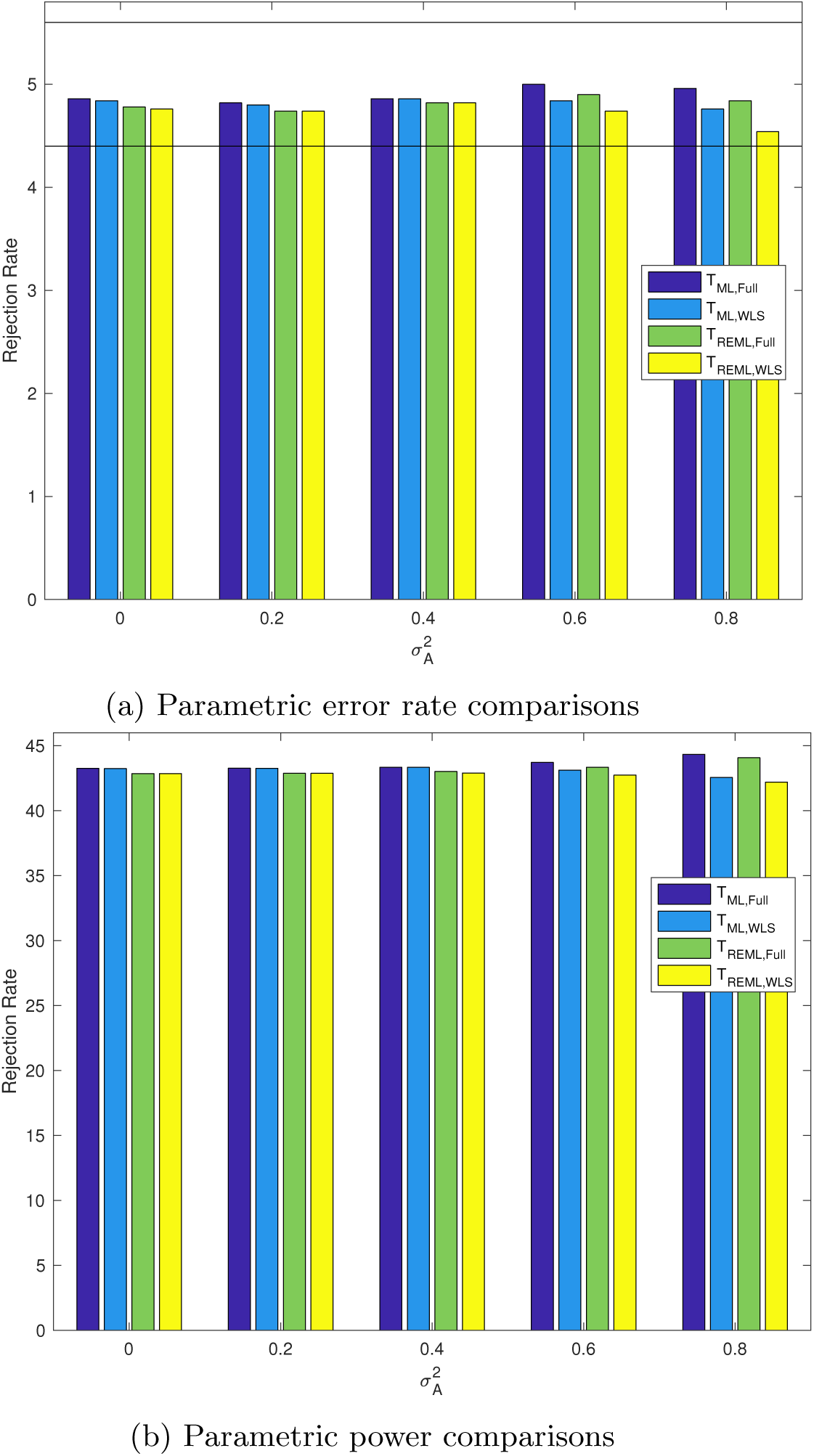
Simulation 2, comparing the simplified ML and REML score test parametric rejection rates using the one-step and the fully converged random effect estimator, 5% nominal (a) and power (b) based on simulation using either a GRM from 300 unrelated individuals or a kinship from GOBS study with 171 individuals and 10 families and 5000 realizations. Monte Carlo confidence interval is (4.40%, 5.60%). Regardless of kinship matrix in the simulation and variance component estimator, non-iterative or fully converged, all methods have similar performances.

**Figure 4:**
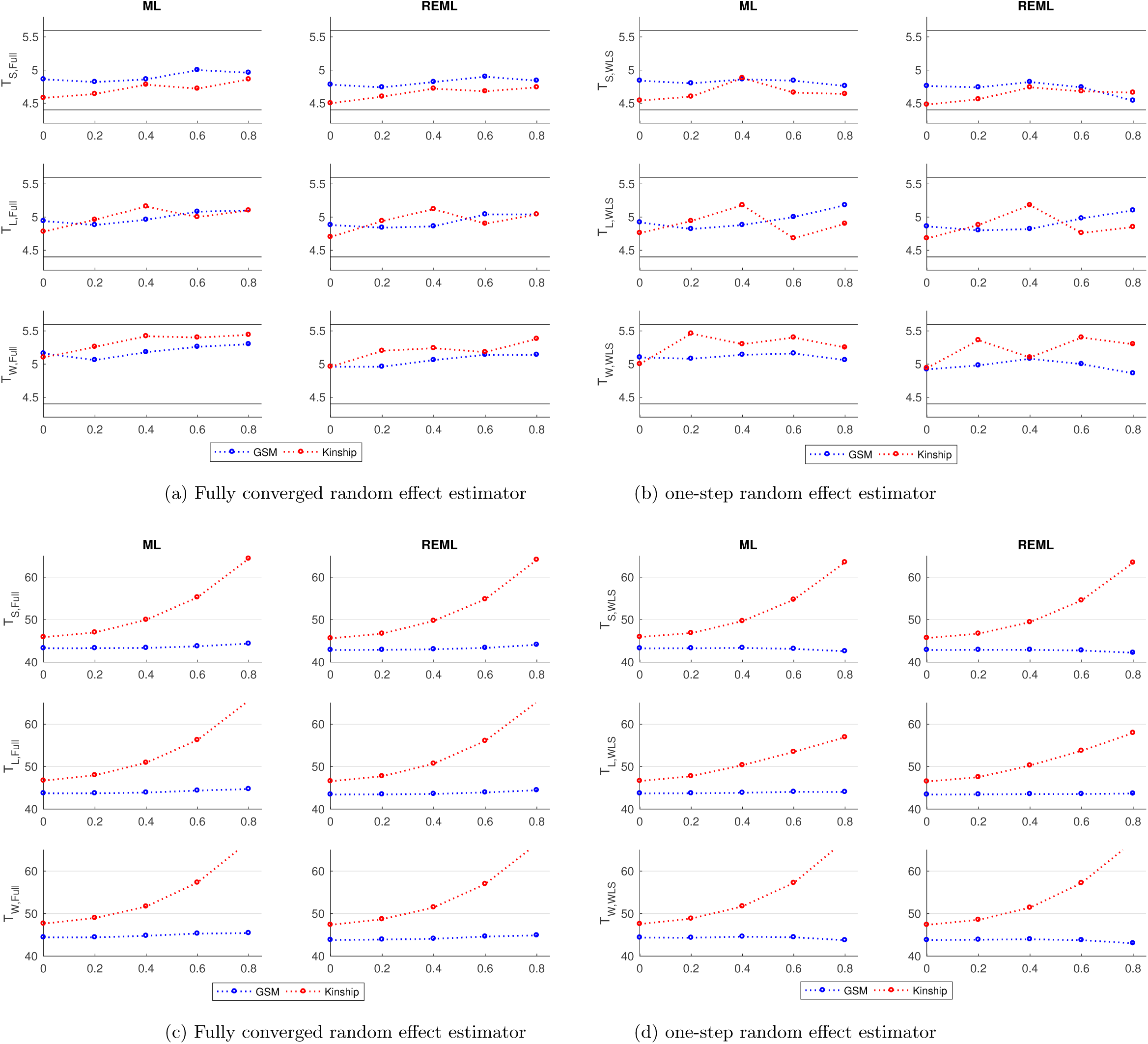
Simulation 2, comparing proposed statistics parametric error rates, 5% nominal (Top panels) and power (bottom panels) based on simulation using either a GRM from 300 unrelated individuals or a kinship from GOBS study with 171 individuals and 10 families and 5000 realisations. The panels (a) and (c) correspond to association statistics using the fully converged random effect estimator and (b) and (d) show the result using the non-iterative random effect estimator. Monte Carlo confidence interval is (4.40%, 5.60%). Regardless of kinship matrix in the simulation and variance component estimator, non-iterative or fully converged, all statistic have similar performances.

**Figure 5:**
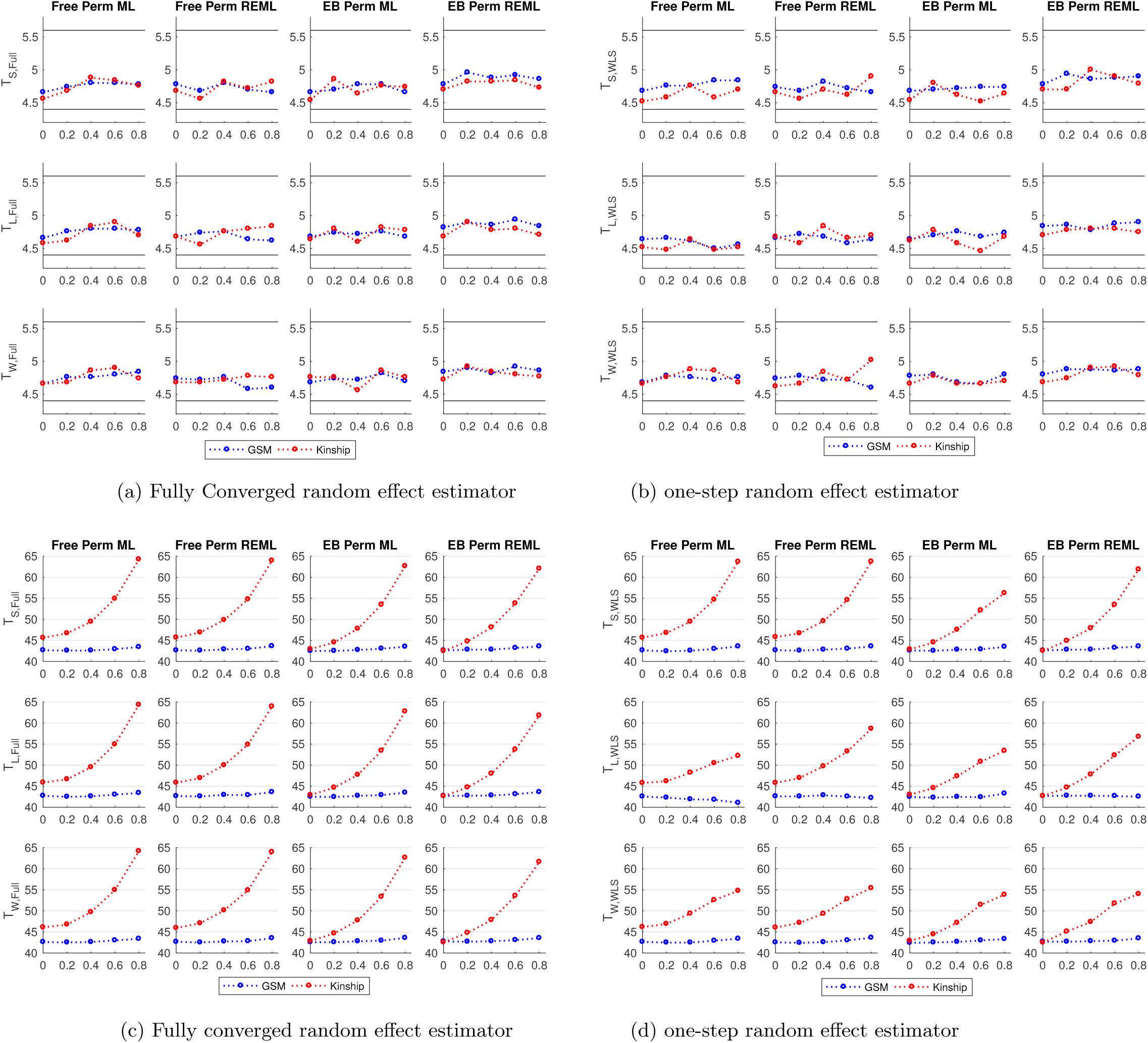
Simulation 2, comparing proposed statistics permutation based error rates, 5% nominal (Top panels) and power (Bottom panels) based on simulation using either a GRM from 300 unrelated individuals or a kinship from GOBS study with 171 individuals and 10 families and 5000 realisations and 500 permutations each realisations. The panels (a) and (c) correspond to association statistics using the fully converged random effect estimator and (b) and (d) show the result using the non-iterative random effect estimator. Monte Carlo confidence interval is (4.40%, 5.60%). Despite the kinship matrix in the simulation and variance component estimator, non-iterative or fully converged, all statistic have similar performances. Both permutation schemes could control the error rate at the nominal level, however free permutation is slightly more powerful than the restricted permutation.

**Figure 6:**
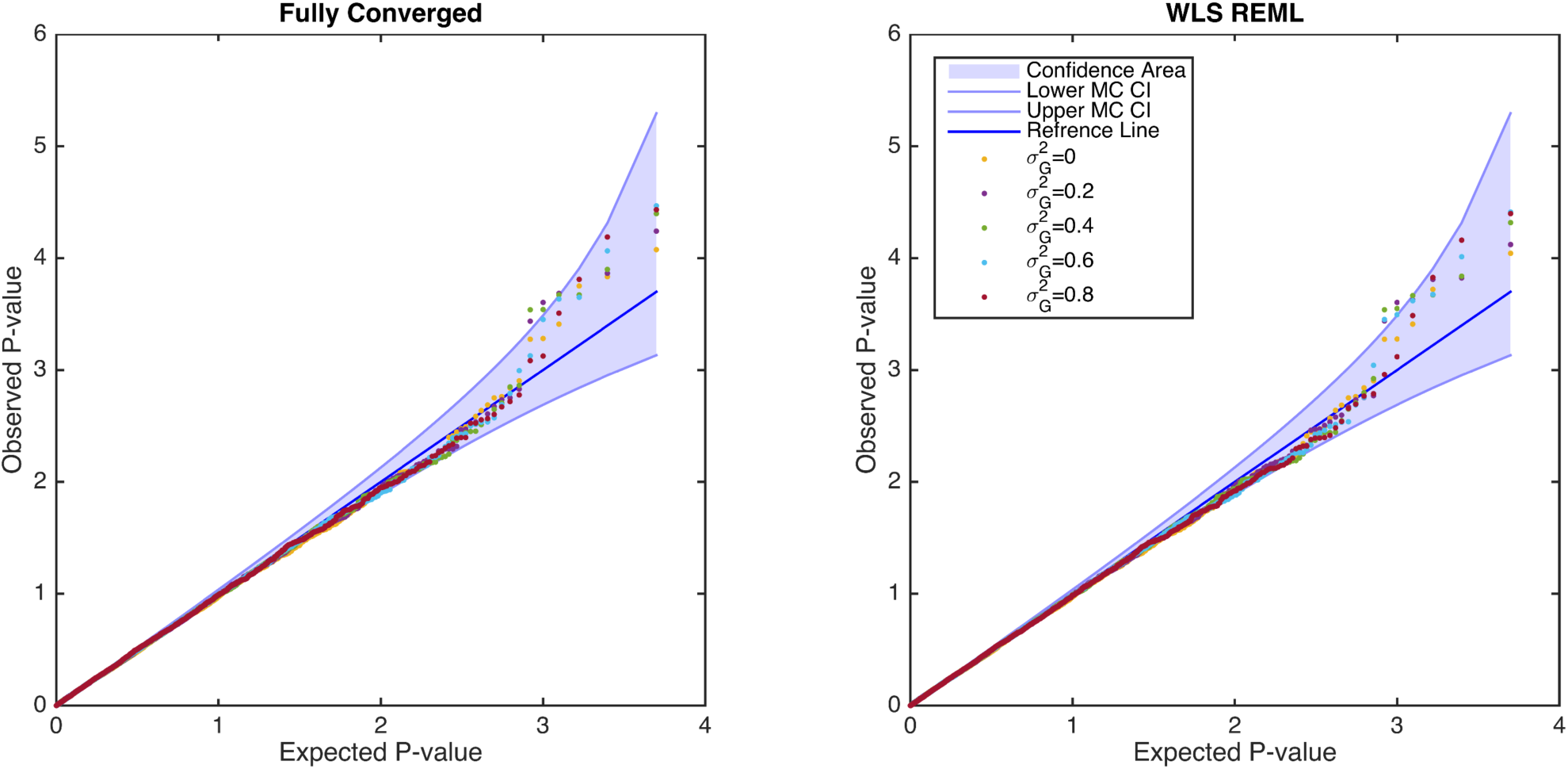
Simulation 3, comparing score statistic parametric null distribution for *H*_0_: *β*_1_ = 0 derived from the simplified REML function using non-iterative and fully converged random effect estimator, for a kinship from GOBS study using 10 families and 171 individuals. There is no apparent difference between the two random effect estimators, and both are consistent with a valid (uniform) P-value distribution.

**Figure 7:**
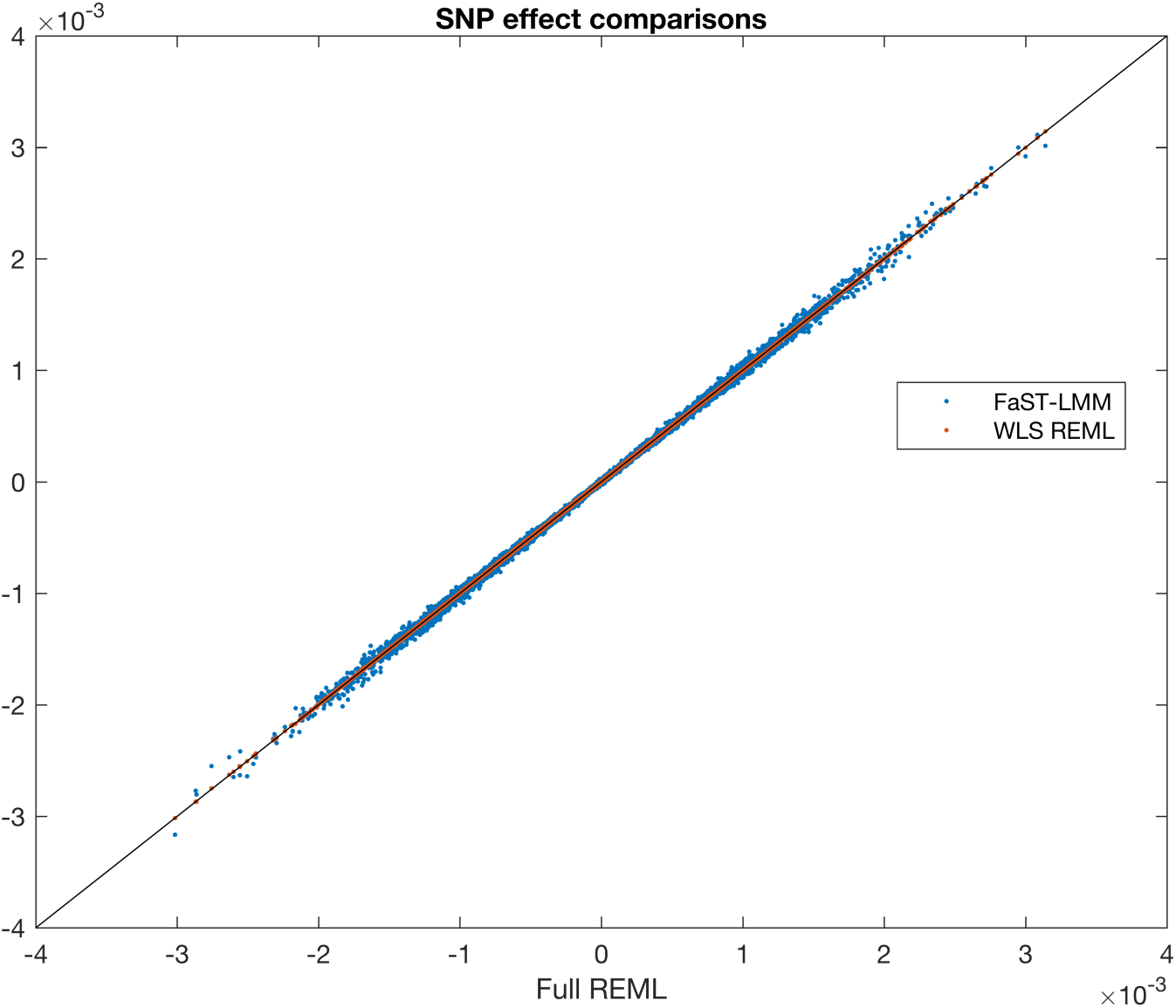
Simulation 4, comparing fixed effect estimation bias between FaST-LMM and the simplified REML function. Each point represents a simulated SNP bias where 6000 SNPs with MAF range (0.05,0.5) are simulated. The x-axis shows parameter estimates bias (*β*_1_) from Full REML and y-axis represents parameter estimates bias over 5000 phenotype realisations using FaST-LMM (blue points) and WLS-REML (red points).

**Figure 8:**
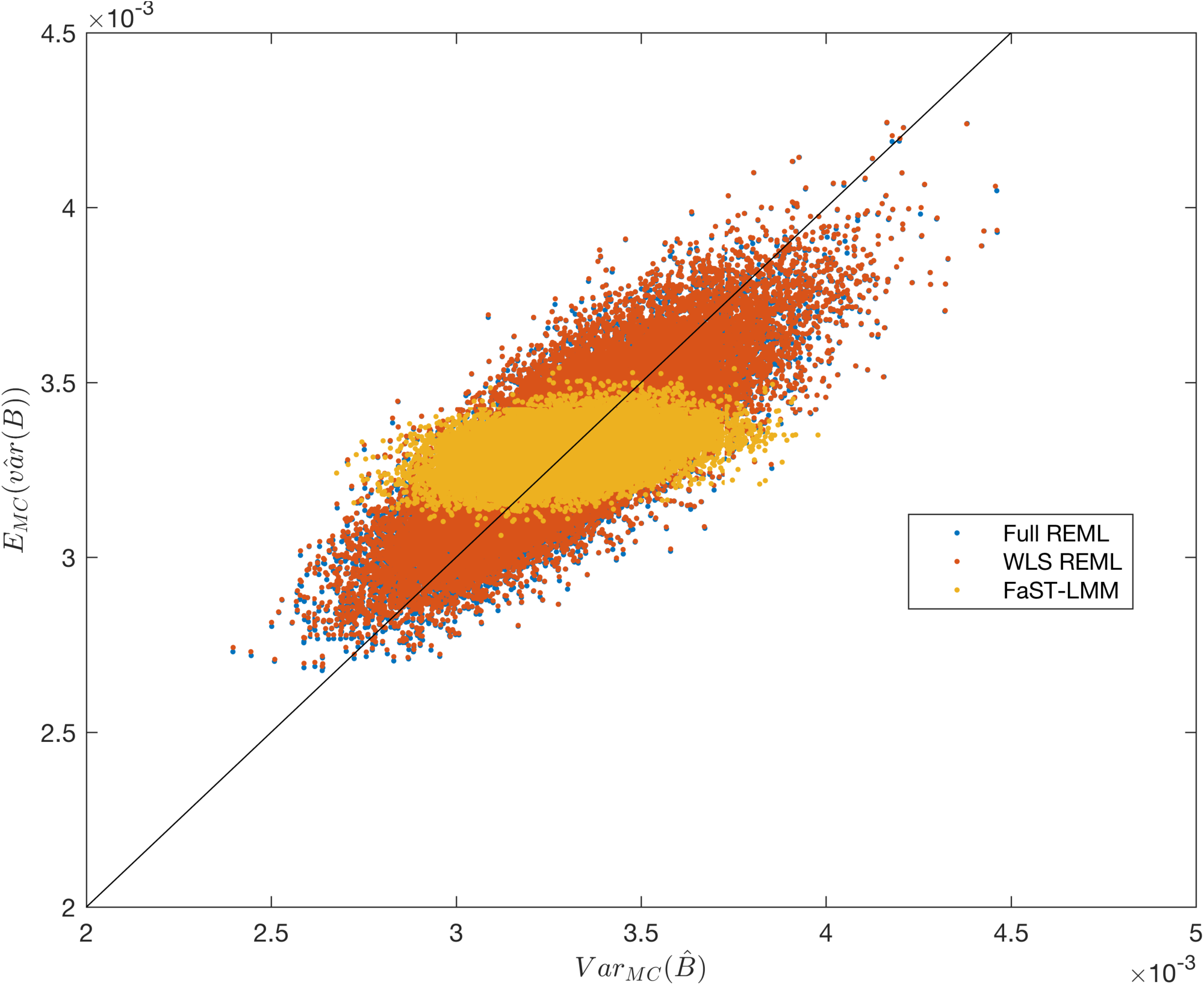
Simulation 4, comparing the Monte Carlo (x-axis) and method-estimated (y-axis) variance of the fixed effect estimator, for FaST-LMM and the simplified REML function. There are 6000 points, one for each simulated SNP, MAF ranging from 0.05 to 0.5. For each SNP, 5000 realizations of data for 300 unrelated subjects are generated with *β*_1_ = 0. The Monte Carlo estimate of variance is over the 5000 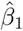; each method’s produces one 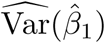 for each realization, which are averaged to obtain the y-axis value.

**Figure 9:**
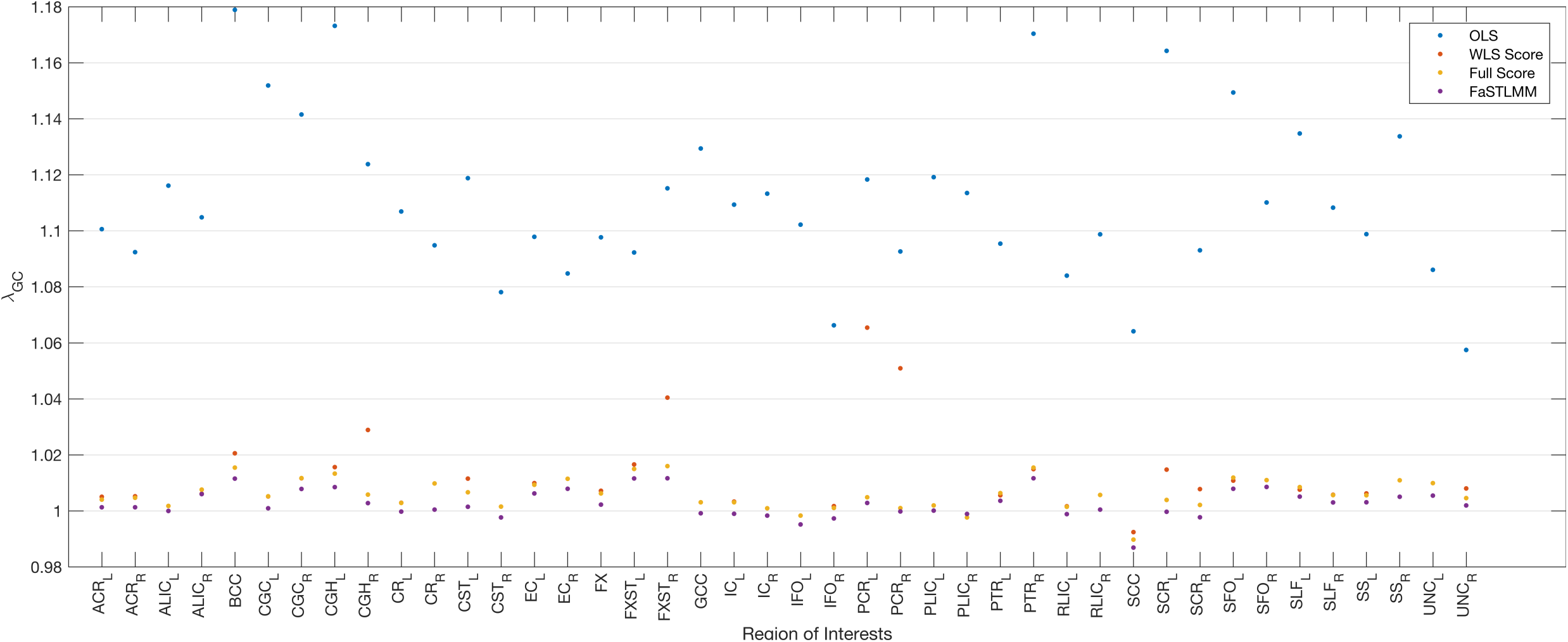
Real data analysis, comparing the genomic control values of the score test based on the simplified REML function using either fully converged random effect estimator (Fully Score, yellow dots) or the WLS-REML random effect estimator (WLS Score, red dots), FaSTLMM (purple dots) with the linear regression with MDS as nuisance fixed effects (OLS, blue dots) for 42 ROIs in the CEU sample. Our proposed method consistently gives smaller genomic factor regardless of random effect estimation method and close to FaST-LMM.

**Figure 10:**
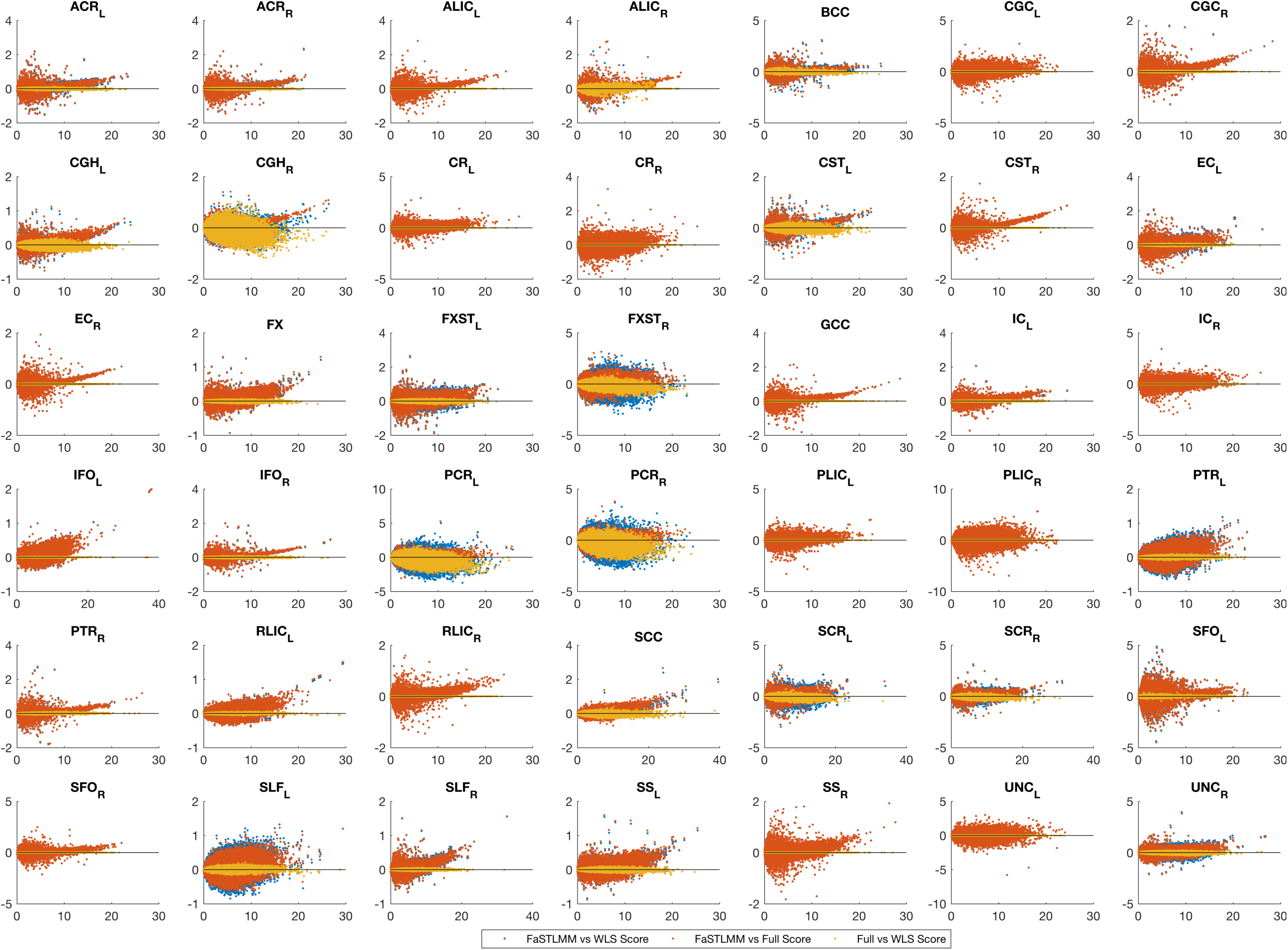
Real data analysis, Bland-Altman plot for comparing values of FaST-LMM and the score test for association testing (*H*_0_: *β*_1_ = 0) using non-iterative and fully converged random effect estimators. Each plot represents a ROI where x-axis shows average statistics and y-axis represents differences. The score tests are almost identical and slightly less powerful than FaST-LMM.

**Figure 11:**
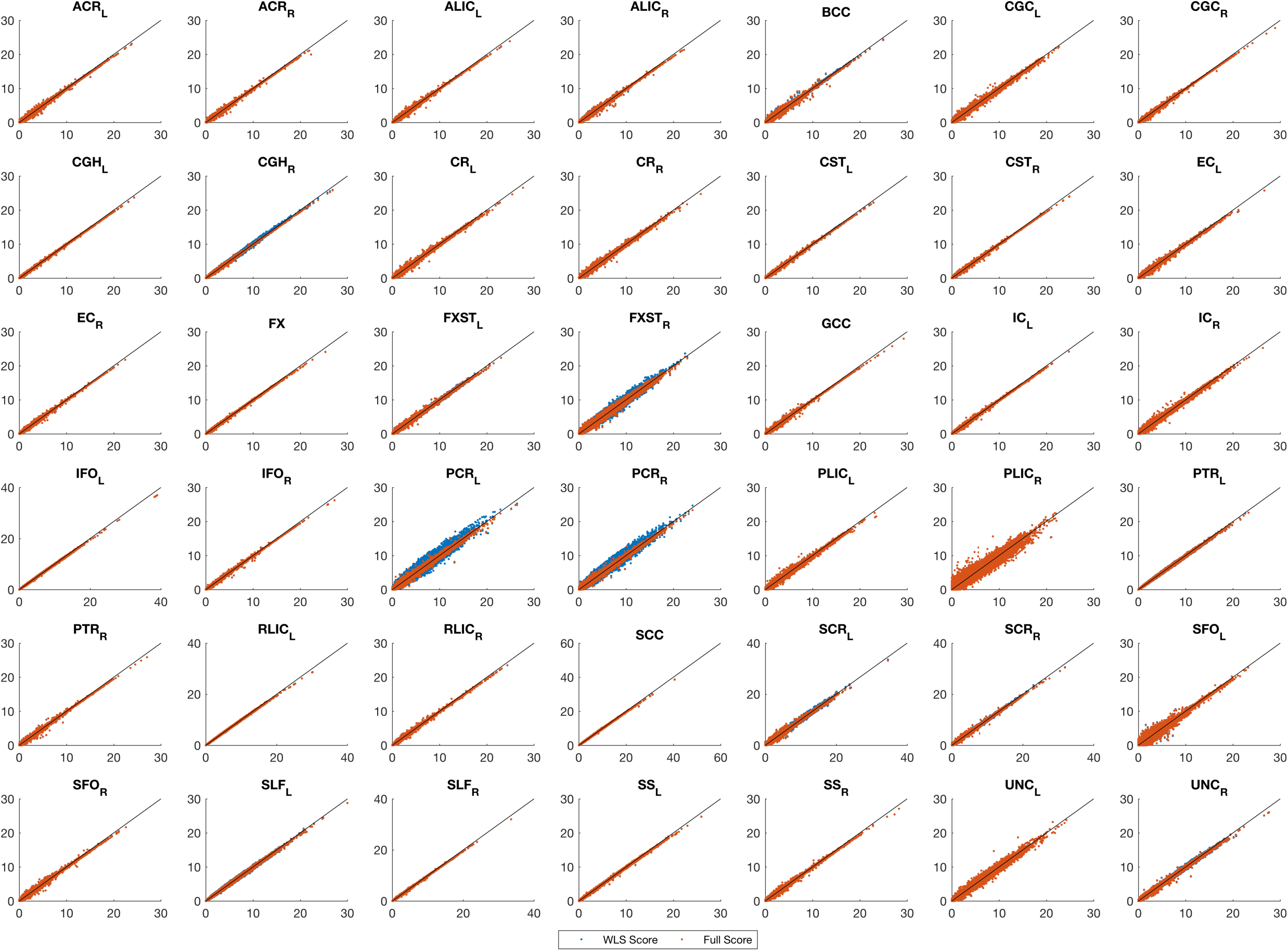
Real data analysis, comparing values of the score test for association testing (*H*_0_: *β*_1_ = 0) using non-iterative and fully converged random effect estimators. and Fast-LMM Each plot represents a ROI where x-axis shows FaST-LMM LRT values score test using estimator and y-axis represents score test using either WLS-REML or the fully converged random effect estimator. The score tests are almost identical and slightly less powerful than FaST-LMM.

**Figure 12:**
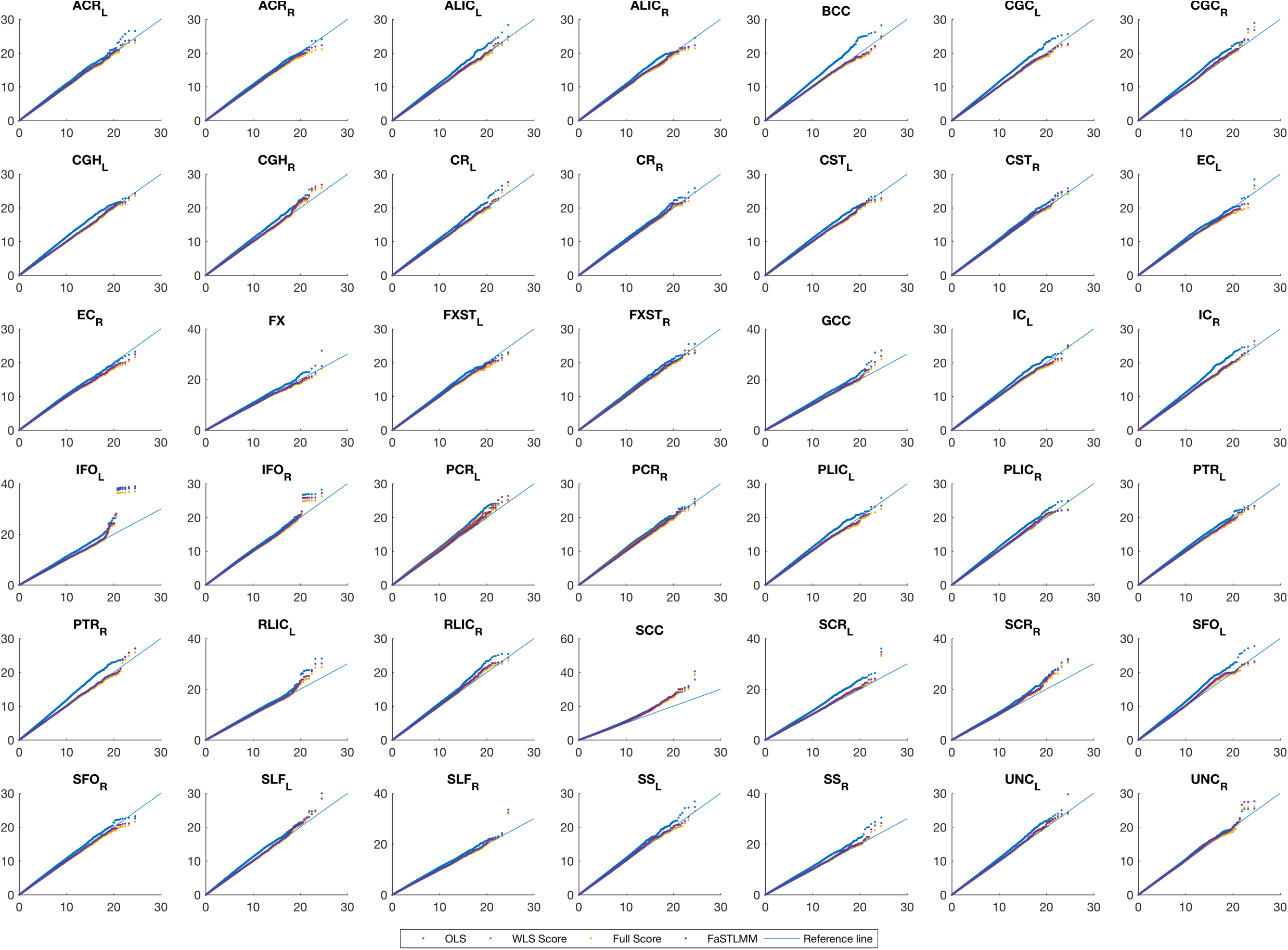
Real data analysis, QQ plot for comparing FaST-LMM and the score test based on the simplified REML function using the WLS-REML random effect estimator with the linear regression with MDS as nuisance fixed effects. Each plot corresponds to different ROIs. These plots show either an identical distribution or slightly larger values for the OLS approach. However the OLS approach has poor genomic control (Figure 9).

**Figure 13:**
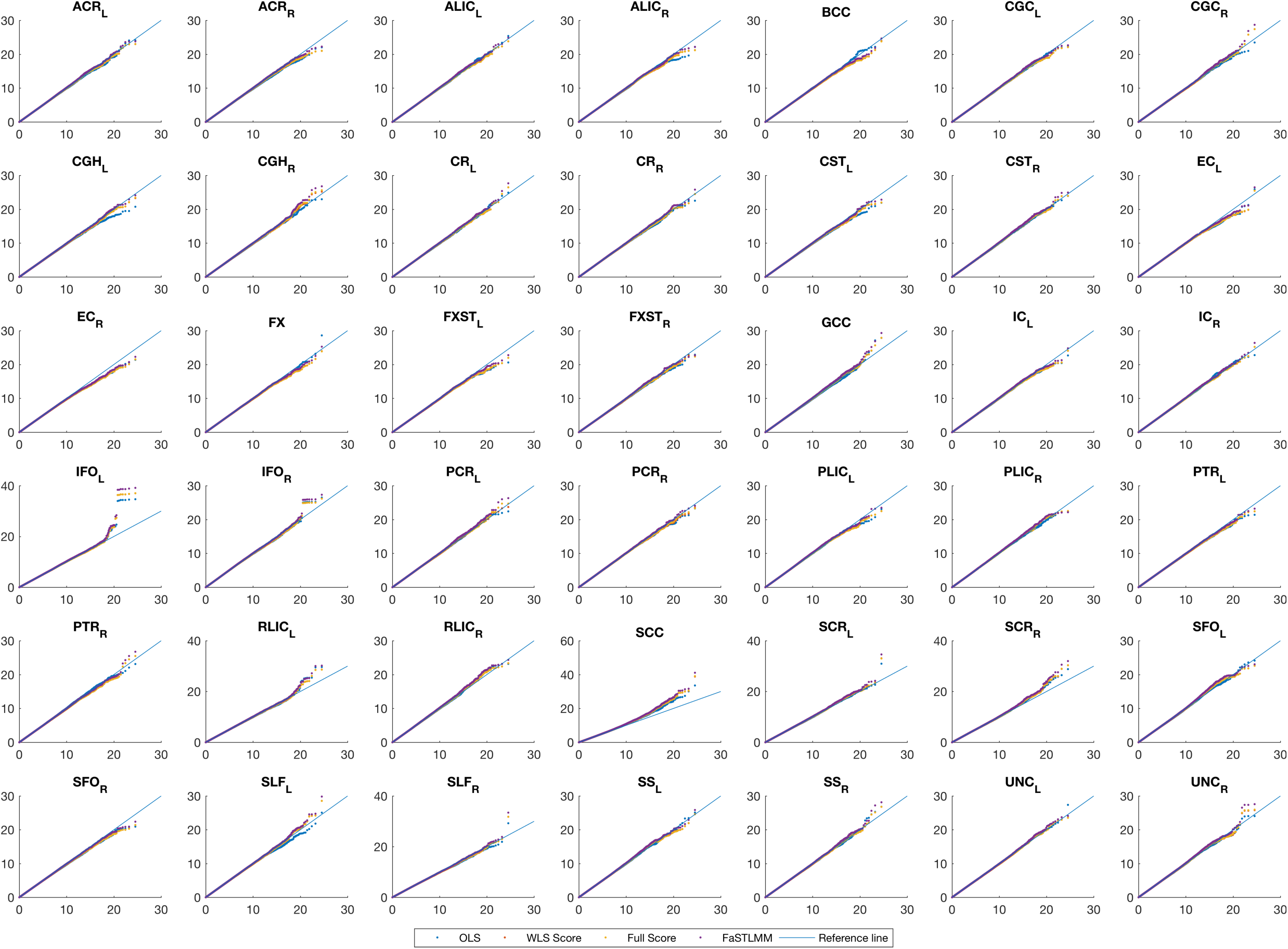
Real data analysis, QQ plot for comparing the adjusted association statistics for genomic control values. Each plot corresponds to different ROIs. These plots show after adjustment we get essentially identical results for the score test based on the simplified REML function using the WELS REML random effect estimator and the OLS approach.

## References

1. Hibar, D. P., Stein, J. L., et al. Franke, B., Thompson, P. M. & Medland, S. E. Common genetic variants influence human subcortical brain structures. Nat. 520, 224–229 (2015). DOI http://dx.doi.org/10.1038/nature14101.

2. Stein, J. L., Medland, S. E., Vasquez, A. A., Hibar, D. P. & et al. Thompson P. M, E. N. I. G. t. M.-A. C. Identification of common variants associated with human hippocampal and intracranial volumes. Nat Genet. 44, 552–561 (2012). DOI 10.1038/ng.2250.

3. Stein, J. L. et al. Genome-wide analysis reveals novel genes influencing temporal lobe structure with relevance to neurodegeneration in alzheimer’s disease. NeuroImage 51, 542 – 554 (2010). DOI http://dx.doi.org/10.1016/j.neuroimage.2010.02.068.

4. Stein, J. L. et al. Voxelwise genome-wide association study (vgwas). NeuroImage 53, 1160 – 1174 (2010). DOI http://dx.doi.org/10.1016/j.neuroimage.2010.02.032. ImagingGenetics.

5. Potkin, S. G. et al. Hippocampal atrophy as a quantitative trait in a genome-wide association study identifying novel susceptibility genes for alzheimer’s disease. PLoS ONE 4, 1–15 (2009). DOI 10.1371/journal.pone.0006501.

6. Potkin, S. G. et al. A genome-wide association study of schizophrenia using brain activation as a quantitative phenotype. Schizophr. Bull. 35, 96–108 (2009). DOI 10.1093/schbul/sbn155.

7. Voight, B. F. & Pritchard, J. K. Confounding from cryptic relatedness in case-control association studies. PLoS Genet. 1 (2005). DOI 10.1371/journal.pgen.0010032.

8. Weir, B. S., Anderson, A. D. & Hepler, A. B. Genetic relatedness analysis: modern data and new challenges. Nat. Rev. Genet. 7, 771–780 (2006). DOI doi:10.1038/nrg1960.

9. Pritchard, J. K., Stephens, M., Rosenberg, N. A. & Donnelly, P. Association mapping in structured populations. The Am. J. Hum. Genet. 67, 170 – 181 (2000). DOI http://dx.doi.org/10.1086/302959.

10. Cardon, L. R. & Palmer, L. J. Population stratification and spurious allelic association. The Lancet 361, 598 – 604 (2003). DOI http://dx.doi.org/10.1016/S0140-6736(03)12520-2.

11. Helgason, A., Yngvadóttir, B., Hrafnkelsson, B., Gulcher, J. & Stefánsson, K. An icelandic example of the impact of population structure on association studies. Nat. genetics 37, 90–95 (2005). DOI doi:10.1038/ng1492.

12. Balding, D. A tutorial on statistical methods for population association studies. Nat Rev Genet. 7, 781–791 (2006). DOI 10.1038/nrg1916.

13. Price, A. L., Zaitlen, N. A., Reich, D. & Patterson, N. New approaches to population stratification in genome-wide association studies. Nat. Rev. 11, 459–463 (2010). DOI 10.1038/nrg2813.

14. Price, A. L. et al. Principal components analysis corrects for stratification in genome-wide association studies. Nat. genetics 38, 904–909 (2006). DOI doi:10.1038/ng1847.

15. Lange, K., Westlake, J. & Spence, M. A. Extensions to pedigree analysis iii. variance components by the scoring method. Annals Hum. Genet. 39, 485–491 (1976). DOI 10.1111/j.1469-1809.1976.tb00156.x.

16. Hopper, J. L. & Mathews, J. D. Extensions to multivariate normal models for pedigree analysis. Annals Hum. Genet. 46, 373–383 (1982). DOI 10.1111/j.1469-1809.1982.tb01588.x.

17. Hasstedt, S. J. A mixed-model likelihood approximation on large pedigrees. Comput. Biomed. Res. 15, 295 – 307 (1982). DOI http://dx.doi.org/10.1016/0010-4809(82)90064-7.

18. Boerwinkle, E., Chakraborty, R. & Sing, C. F. The use of measured genotype information in the analysis of quantitative phenotypes in man. Annals Hum. Genet. 50, 181–194 (1986). DOI 10.1111/j.1469-1809.1986.tb01037.x.

19. Almasy, L. & Blangero, J. Multipoint quantitative-trait linkage analysis in general pedigrees. The Am. J. Hum. Genet. 62, 1198–1211 (1998). DOI 10.1086/301844.

20. Yu, J. et al. A unified mixed-model method for association mapping that accounts for multiple levels of relatedness. Nat. genetics 38, 203–8 (2006). DOI 10.1038/ng1702.

21. Kang, H. M. et al. Efficient control of population structure in model organism association mapping. Genet. 178, 1709–1723 (2008). DOI 10.1534/genetics.107.080101.

22. Zhang, Z. et al. Mixed linear model approach adapted for genome-wide association studies. Nat. genetics 42, 355–360 (2010). DOI 10.1038/ng.546.

23. Kang, H. M. et al. Variance component model to account for sample structure in genome-wide association studies. Nat. genetics 42, 348–54 (2010). DOI 10.1038/ng.548.

24. Lippert, C. et al. FaST linear mixed models for genome-wide association studies. Nat. Methods 8, 833–837 (2011). DOI 10.1038/nmeth.1681.

25. Lippert, C. et al. Improved linear mixed models for genome-wide association studies. Nat. methods 8, 833–5 (2011). DOI 10.1038/nmeth.2037.

26. Zhou, X. & Stephens, M. Genome-wide efficient mixed-model analysis for association studies. Nat. genetics 44, 821–4 (2012). DOI 10.1038/ng.2310.

27. Svishcheva, G. R., Axenovich, T. I., Belonogova, N. M., van Duijn, C. M. & Aulchenko, Y. S. Rapid variance components–based method for whole-genome association analysis. Nat. Genet. 44, 1166–1170 (2012). DOI 10.1038/ng.2410.

28. Pirinen, M., Donnelly, P. & Spencer, C. C. A. Efficient computation with a linear mixed model on large-scale data sets with applications to genetic studies. Ann. Appl. Stat. 7, 369–390 (2013). DOI 10.1214/12-AOAS586.

29. Listgarten, J., Lippert, C. & Heckerman, D. FaST-LMM-Select for addressing confounding from spatial structure and rare variants. Nat. Genet. 45, 470–471 (2013). DOI 10.1038/ng.2620.

30. Widmer, C. et al. Further improvements to linear mixed models for genome-wide association studies. Sci. reports 4, 6874 (2014). DOI 10.1038/srep06874.

31. Kadri, N. K., Guldbrandtsen, B., Sørensen, P. & Sahana, G. Comparison of genome-wide association methods in analyses of admixed populations with complex familial relationships. PLoS ONE 9, e88926 (2014). DOI 10.1371/journal.pone.0088926.

32. Yang, J., Zaitlen, N. A., Goddard, M. E., Visscher, P. M. & Price, A. L. Advantages and pitfalls in the application of mixed-model association methods. Nat. genetics 46, 100–6 (2014). DOI 10.1038/ng.2876.

33. Blangero, J. et al. A kernel of truth: statistical advances in polygenic variance component models for complex human pedigrees., vol. 81 (2013).

34. Friston, K. J., Worsley, K. J., Frackowiak, R. S. J., Mazziotta, J. C. & Evans, A. C. Assessing the significance of focal activations using their spatial extent. Hum. Brain Mapp. 1, 210–220 (1994). DOI 10.1002/hbm.460010306.

35. Smith, S. & Nichols, T. Threshold-free cluster enhancement: addressing problems of smoothing, threshold dependence and localisation in cluster inference. Neuroimage 44, 83–98 (2009). DOI 10.1016/j.neuroimage.2008.03.061.

36. Eklund, A., Nichols, T. E. & Knutsson, H. Cluster failure: Why fmri inferences for spatial extent have inflated false-positive rates. Proc. Natl. Acad. Sci. 113, 7900–7905 (2016). DOI 10.1073/pnas.1602413113.

37. Ge, T., Feng, J., Hibar, D. P., Thompson, P. M. & Nichols, T. E. Increasing power for voxel-wise genome-wide association studies: The random field theory, least square kernel machines and fast permutation procedures. NeuroImage 63, 858 – 873 (2012). DOI https://doi.org/10.1016/j.neuroimage.2012.07.012.

38. Nichols, T. & Hayasaka, S. Controlling the familywise error rate in functional neuroimaging: a comparative review. Stat. Methods Med. Res. 12, 419–446 (2003). DOI 10.1191/0962280203sm341ra.

39. Nichols., T. E. & Holmes, A. P. Nonparametric permutation tests for functional neuroimaging: A primer with examples. Hum. Brain Mapp. 15, 1–25 (2002). DOI 10.1002/hbm.1058.

40. Ganjgahi, H. et al. Fast and powerful heritability inference for family-based neuroimaging studies. NeuroImage 115, 256 – 268 (2015). DOI http://dx.doi.org/10.1016/j.neuroimage.2015.03.005.

41. Clopper, C. J. & Pearson, E. S. The use of confidence or fiducial limits illustrated in the case of the binomial. Biom. 26, 404–413 (1934). DOI 10.2307/2331986.

42. Amemiya, T. A note on a heteroscedastic model. J. Econom. 6, 365 – 370 (1977). DOI http://dx.doi.org/10.1016/0304-4076(77)90006-9.

43. Rao, C. R. Linear Statistical Inference and its Applications (John Wiley & Sons, Inc., 2008).

44. Neyman, J. & Pearson, E. S. On the problem of the most efficient tests of statistical hypotheses. Philos. Transactions Royal Soc. Lond. Ser. A, Containing Pap. a Math. or Phys. Character 231, pp. 289–337 (1933).

45. Winkler, A. M., Ridgway, G. R., Webster, M. A., Smith, S. M. & Nichols, T. E. Permutation inference for the general linear model. NeuroImage 92C, 381–397 (2014). DOI 10.1016/j.neuroimage.2014.01.060.

46. Abney, M. Permutation testing in the presence of polygenic variation. Genet. Epidemiol. 39, 249–258 (2015). DOI 10.1002/gepi.21893.

47. Cheng, R. & Palmer, A. A. A simulation study of permutation, bootstrap, and gene dropping for assessing statistical significance in the case of unequal relatedness. Genet. 193, 1015–1018 (2013). DOI 10.1534/genetics.112.146332.

## References

D B Allison, Michael C. Neale, R Zannolli, N J Schork, C I Amos, and J Blangero. Testing the robustness of the likelihood-ratio test in a variance-component quantitative-trait loci-mapping procedure. American journal of human genetics, 65(2):531–44, 1999. doi: http://dx.doi.org/10.1086/302487.

Neda Jahanshad, Peter V. Kochunov, Emma Sprooten, Ren C. Mandl, Thomas E. Nichols, Laura Almasy, John Blangero, Rachel M. Brouwer, Joanne E. Curran, Greig I. de Zubicaray, Ravi Duggirala, Peter T. Fox, L. Elliot Hong, Bennett A. Landman, Nicholas G. Martin, Katie L. McMahon, Sarah E. Medland, Braxton D. Mitchell, Rene L. Olvera, Charles P. Peterson, John M. Starr, Jessika E. Sussmann, Arthur W. Toga, Joanna M. Wardlaw, Margaret J. Wright, Hilleke E. Hulsho? Pol, Mark E. Bastin, Andrew M. McIntosh, Ian J. Deary, Paul M. Thompson, and David C. Glahn. Multi-site genetic analysis of diffusion images and voxelwise heritability analysis: A pilot project of the enigmadti working group. NeuroImage, 81:455 – 469, 2013. doi: http://dx.doi.org/10.1016/j.neuroimage.2013.04.061.

Hyun Min Kang, Noah A. Zaitlen, Claire M. Wade, Andrew Kirby, David Heckerman, Mark J. Daly, and Eleazar Eskin. Efficient control of population structure in model organism association mapping. Genetics, 178(3):1709–1723, 2008. doi: 10.1534/genetics.107.080101.

Christoph Lippert, Jennifer Listgarten, Ying Liu, Carl M. Kadie, Robert I. Davidson, and David Heckerman. FaST linear mixed models for genome-wide association studies. Nature Methods, 8 (10):833–837, 2011. doi: 10.1038/nmeth.1681.

Shayle R Searle, George Casella, and Charles E McCulloch. Variance components, volume 391. John Wiley & Sons, 2009. doi:10.1002/9780470316856.

Bertrand Servin and Matthew Stephens. Imputation-based analysis of association studies: Candidate regions and quantitative traits. PLoS Genet, 3(7), 07 2007. doi:10.1371/journal.pgen. 0030114.

Stephen M. Smith, Mark Jenkinson, Heidi Johansen-Berg, Daniel Rueckert, Thomas E. Nichols, Clare E. Mackay, Kate E. Watkins, Olga Ciccarelli, M. Zaheer Cader, Paul M. Matthews, and Timothy E.J. Behrens. Tract-based spatial statistics: Voxelwise analysis of multi-subject diflusion data. NeuroImage, 31(4):1487 – 1505, 2006. doi: http://dx.doi.org/10.1016/j.neuroimage.2006.02.024.

